# The growth-promoting protein TMEM263 is an ER resident that controls early stages of lipid droplet biogenesis

**DOI:** 10.1101/2025.07.04.663055

**Authors:** Katy Yalci, Nafsika Chala, Joanna Moss, Przemysław Zakrzewski, Rabia Sevil, Lana Willoughby, Seren Wonklyn, Thomas Andrews, Chrissy Hammond, Robin A. Corey, Bernadette Carroll

## Abstract

Cell and organismal growth is controlled not only by the availability of nutrients, but also the ability to dynamically sense and respond to changes in metabolic demand. We have identified a protein, TMEM263 (also known as C12orf23) as a new mechanistic link between growth and lipid metabolism. TMEM263 was discovered in a screen as an ER-resident protein, and we have characterised its role as both necessary and sufficient for lipid droplet formation. TMEM263 has two transmembrane domains that fold into a hairpin structure which are essential for its localisation to the ER and to lipid droplets. Functionally, we propose that TMEM263 can interact with and support the condensation of neutral lipids in a bilayer to promote lipid droplet formation and growth. Consistently, loss of TMEM263 in cells and in zebrafish significantly impairs lipid droplet formation. Loss of TMEM263 protein function *in vivo* is associated with organismal growth defects and proportional dwarfism and our study provides a mechanistic understanding linking these phenotypes to impaired lipid droplet biology.

## Introduction

The ability to sense and appropriately respond to nutrients is essential for the control of cell and whole-body growth. Too few or too many nutrients is widely associated with human disease and thus it is crucial we understand the molecular determinants controlling nutrient handling and nutrient homeostasis. An often-overlooked regulator of nutrient homeostasis and cellular growth is the endoplasmic reticulum (ER). It orchestrates the synthesis of proteins and lipids, facilitates proper protein folding, and serves as a store of intracellular calcium. To support these activities, the ER hosts a diverse array of resident proteins; ER-resident chaperones, for instance, are critical for the folding and maturation of secretory and membrane proteins, including growth factor receptors that directly influence cell growth and survival^1, 2^. The localisation of such ER-resident proteins is tightly regulated, in part by the heptameric coatomer protein complex (or COPI). COPI recognises specific cargo motifs and facilitates retrograde transport from the cis-Golgi back to the ER to prevent proteins being lost via the secretory pathway, as well as intra-Golgi trafficking^3,4^. COPI is essential for maintaining ER-Golgi homeostasis; its disruption results in widespread defects in vesicular transport, ER and Golgi disorganization, mitochondrial dysfunction, and the aberrant accumulation of autophagosomes and lipid droplets^5-7^.

The ER also plays a central role in the dynamic sensing and response to metabolic challenges, including nutrient starvation and surplus^8^. For example, during nutrient starvation, recruitment of proteins such as WIPI2 and ATG16L to ER subdomains rich in P13KC3-dependent PI3P promotes the formation of phagophores, the precursors of autophagosomes which support nutrient liberation and cell survival by delivering cytoplasmic cargo to the lysosome for degradation. One consequence of this degradation is an increase in free fatty acids from the breakdown of membranes ^9^. Similar fatty acid accumulation can occur as a result of increased *de novo* biosynthesis/decreased utilisation or increased uptake from the environment via cell surface receptors such as CD36 ^10-13^. Fatty acids represent an important fuel source, while excess lipid accumulation can cause lipotoxicity, cytotoxic stress and cell death^10, 11^. Therefore as a protective storage mechanism, fatty acids are converted by ER-resident enzymes into neutral lipids which are stored in lipid droplets. Specifically, enzymes such as diacylglycerol O-acyltransferase 1 (DGAT1) and acyl-CoA:cholesterol acyltransferases (ACAT1/2) esterify fatty acids and cholesterol, respectively, generating triacylglycerols (TAGs) and cholesterol esters which accumulate within the ER phospholipid bilayer. When the neutral lipid accumulation surpasses a critical concentration (∼5–10 mol%), they undergo phase-separation into a lipid ‘lens’. The growing lipid core grows asymmetrically towards the cytoplasm in a process regulated by the biophysical properties of the ER bilayer such as membrane tension and phospholipid composition^10, 14^. The subsequent monolayer-enclosed lipid droplet remains functionally and physically connected to the ER by extensive membrane contact sites. These contact sites facilitate the exchange of proteins and lipids to allow the rapid response to changes in cellular metabolic demand, leading to lipid droplet expansion, shrinkage or degradation ^10-13^.

Several ER-resident proteins have emerged as key regulators of lipid droplet biogenesis, acting in concert with metabolic enzymes. Seipin/BSCL2 is a multimeric complex that localises at ER–lipid droplet contacts sites to coordinate the nucleation and growth of lipid droplets by organising lipid lenses and recruiting metabolic enzymes^15-17^. FIT2, another ER-resident protein, is implicated in early lipid droplet formation, potentially through modulating membrane curvature^18^. Furthermore, several ER-resident integral membrane proteins can transition to the lipid droplet monolayer via the ER-to-LD (ERTOLD) pathway and are referred to as class I lipid droplet proteins^10^. They have diverse functions; for example, perilipin 1 controls access of lipases^19^ and other proteins to the lipid droplet surface, while GPAT4 is a lipid biosynthetic enzyme^20^. Given the tight and dynamic spatial relationship between the two organelles, there are likely to be many more ER-resident proteins involved in lipid droplet biology.

In this study, we used unbiased proteomics to identify ER-resident proteins that control mammalian growth. We identified several proteins associated with lipid metabolism as ER-resident proteins, and characterised one, TMEM263/C12orf23 as a lipid droplet protein. Our results indicate TMEM263 can transition from the ER to lipid droplets, and its overexpression appears to be sufficient to induce the accumulation of enlarged, clustered lipid droplets, particularly upon lipid feeding. Our modelling data indicate that TMEM263 has two transmembrane (TM) domains that fold into a hairpin structure and our experimental data demonstrate that these are essential for its localisation and function. Molecular dynamics simulations indicate that TMEM263 can interact with and support the condensation of neutral lipids in a bilayer, and mutation of these lipid-interacting residues impairs TMEM263-induced lipid droplet formation in cells. Consistently, loss of TMEM263 in cells and in zebrafish, significantly reduces lipid droplet numbers following lipid feeding. Together, our data indicate TMEM263 is required for efficient lipid droplet formation *in vitro* and *in vivo and* provides a mechanistic understanding for the observed organismal growth defects and proportional dwarfism observed upon TMEM263 loss*-*of-function^21-25^.

## Results

The coatomer, or COPI protein complex is essential for the maintenance of cell homeostasis and growth. It is a large stable protein complex of seven subunits (α, β, β’, γ, δ, ε, ζ) that is recruited to cis-Golgi membranes via Arf1-GTP to promote vesicle biogenesis and budding, to facilitate cargo transport either within the Golgi or back to the ER. Cargo proteins interact with outer-COPI coat subunits which consists of α-COP, β’-COP and ε-COP predominantly via the short consensus motifs, KxKxx or KKxx present in the very C-terminal of their cytoplasmic tails^26^. Additional, less well-characterised motifs including RxR and Φ-(K/R)-X-L-X-(K/R), are recognised by β-COPI / δ-COPI and δ-COPI / ζ-COPI, respectively^4^. To identify new ER-resident COPI cargo which are important for controlling cell growth, we overexpressed a component of the outer-coat COPI complex, GFP-ε-COPI in HeLa cells and initially confirmed it integrated properly into complex with other COPI machinery and localised to the ER and Golgi **(Fig.S1A,B)**. Subsequent GFP immunoprecipitation and proteomics analyses identified 473 high confidence (log FC ≥2.5, p-value ≤0.005) binding partners for COPI, including 105 that contain the canonical COP-I binding motifs, KxKxx or KKxx, or RxR **(Fig.1A, S1C)**. We further filtered the results using the online MGI database^27, 28^ to identify proteins that have been implicated in mammalian growth. This approach produced a list of 22 proteins, several of which are also implicated in lipid metabolism **(Fig.1A, S1C)**. These include UGT8 which is a ceramide galactosyltransferase and HSD17B12 which is a 17-beta-hydroxysteroid dehydrogenase. Other interesting hits include TMEM41B which is a lipid scramblase that has been implicated in the biogenesis of lipid droplets and autophagosomes from the ER membrane^29, 30^, and TMEM263. Also known as C12orf23, TMEM263 is a vertebrate-specific 12kDa protein with a poorly characterised structure and function. Loss-of-function mutations in TMEM263 have been associated with bone and growth defects in chickens, mice and humans^21-25^. Mechanistically, human GWAS studies implicate *Tmem263* variants in bone mineral density and osteoblast function^21, 22^. The defects in bone growth observed in *Tmem263* knock-out mice were associated with reduced growth hormone receptor expression, and a subsequent reduction in circulating IGF-1 levels^23^, although there were no observed differences in circulating IGF-1 in mutant chicken strains^24, 31, 32^. Thus, it is unclear at present whether these phenotypes were a direct result of TMEM263 loss, and the molecular function of TMEM263 remains largely unknown. Interestingly, TMEM263 has also been validated as a high confidence, but uncharacterised lipid droplet protein^12, 33^.

**Figure 1.**
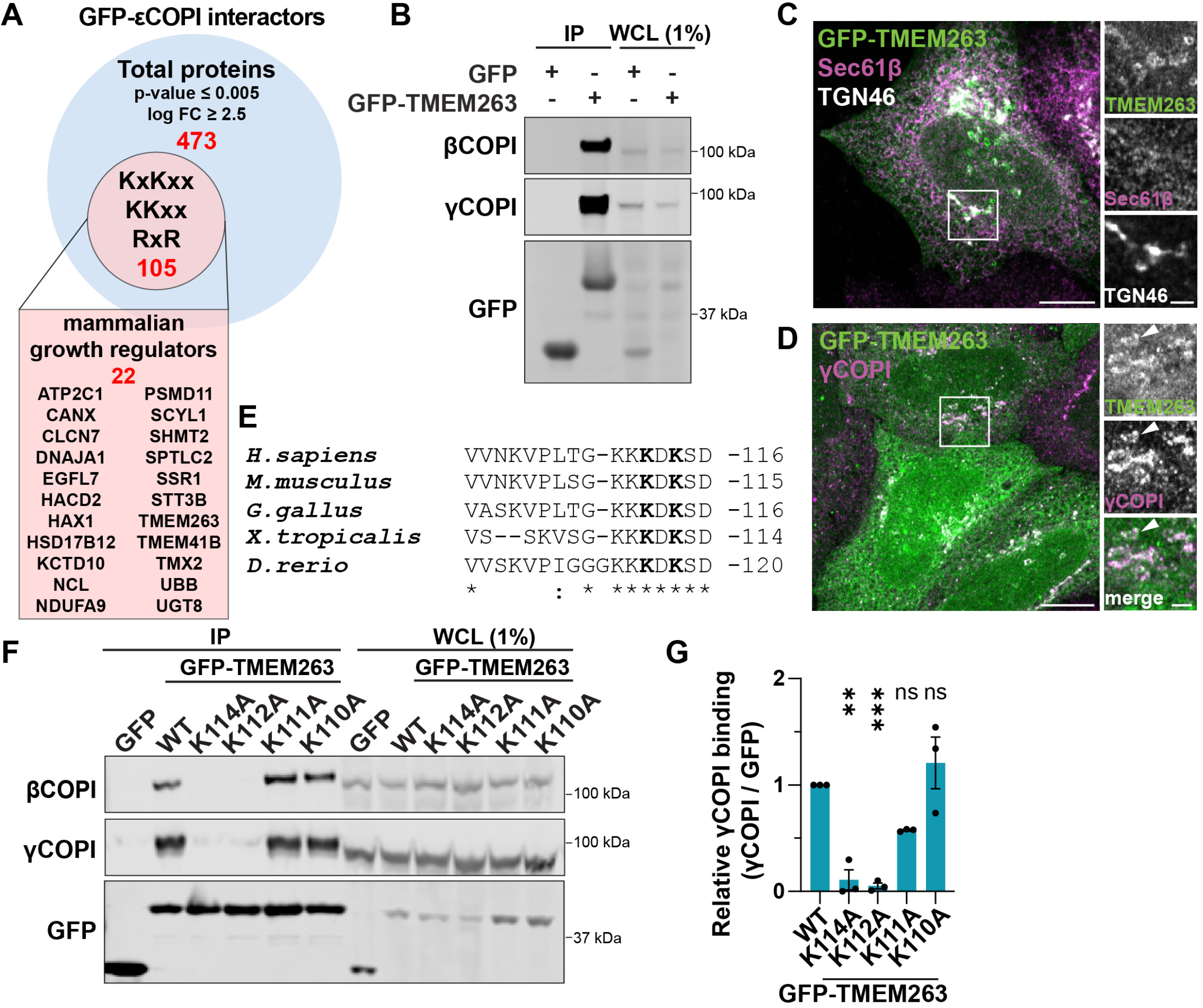
TMEM263 is a COPI cargo protein. **(A)** Identification of COPI cargo. HeLa cells stably expressing GFP or GFP-εCOPI were subject to GFP immunoprecipitation and analysis by TMT mass spectrometry. COPI cargoes were defined as proteins with a log fold change (FC) ≥ 2.5 and p-value ≤0.005 GFP-εCOPI / GFP (n = 4). Proteins were further filtered for those containing COPI-interacting motifs, either KxKxx or KKxx (at the C-terminus) or RxR (within the last 30 amino acids of C-terminus), and associated with mammalian growth regulation. **(B)** TMEM263 is a COPI-interacting protein. HeLa cells expressing GFP-TMEM263 were subject to GFP immunoprecipitation and western blot to detect endogenous COPI components, γCOPI and βCOPI. **(C,D)** TMEM263 localisation is consistent with COPI cargo. HeLa cells expressing GFP-TMEM263 were fixed and immunostained with antibodies against endogenous Sec61β (ER) and TGN46 (trans-Golgi) **(C)** or γCOPI **(D). (E)** TMEM263 contains a canonical COPI interacting motif. ClustalW alignment highlighting a conserved canonical COPI motif, KxKxx in the terminal five amino acids of TMEM263. **(F,G)** Mutation of COPI-binding motif inhibits TMEM263-COPI interaction. HEK293T cells overexpressing GFP-TMEM263 wild-type or mutants, as indicated were subject to GFP immunoprecipitation and western blot to detect endogenous COPI components, γCOPI and βCOPI **(F)**. The interaction between GFP-TMEM263 and γCOPI was quantified **(G)**. Unless stated otherwise, n=3 and error bars represent Standard Error of the Mean (SEM). Ordinary one-way ANOVA with Dunnett’s multiple comparisons test was performed. Scale bars: 10 μm and 2 μm for zoomed images. ns p ≥ 0.05, ** p < 0.01, *** p < 0.001

To confirm that TMEM263 is an ER-resident protein and COPI cargo, we validated the COPI-TMEM263 interaction by immunoprecipitation using GFP-ε–COPI and mCherry-TMEM263 **(Fig.S1D)**. The reciprocal immunoprecipitation using GFP-TMEM263 as bait further confirmed a robust interaction with endogenous COPI **(Fig.1B)**. As expected, GFP-TMEM263 colocalised with the ER, Golgi (**Fig.1C)** and endogenous COPI **(Fig.1D)**, the latter at small vesicular structures and at a region consistent with the Golgi. The C-terminal five amino acids of TMEM263 encode the canonical COPI binding motif, KxKxx (**K**D**K**SD) **(Fig.1E)** and mutation of lysines in this motif (amino acids 112 and 114) abolished the interaction with COPI **(Fig.1F,G)**. As a control, mutation of two upstream lysines residues (K110 and K111) had no significant impact on TMEM263-COPI binding **(Fig.1F,G)**. These data validate TMEM263 a bona fide COPI cargo protein.

We observed that upon overexpression, GFP-TMEM263 showed a heterogenous distribution, with a clear localisation to regions and structures of the cell consistent with the ER and Golgi, but also at vesicular structures throughout the cytoplasm **(Fig.1C,D and Fig.2A)**. GFP-TMEM263 positive puncta were observed in ∼80% transfected HeLa cells, which ranged in size from tiny punctate structures to large, hollow vesicles **(Fig.2A, Fig.S2A)**. The latter structures resembled lipid droplets, and to confirm this, we stained cells with the neutral lipid dye, and lipid droplet marker, LipidTOX. TMEM263-positive puncta were almost exclusively positive for the LipidTOX **(Fig.S2B)**. Overexpression of GFP-TMEM263 showed a significant increase in LipidTOX-positive area in both vehicle control (BSA) and oleic acid (OA)-fed HeLa cells, above the signal in neighbouring, untransfected controls **(Fig.2B-E, Fig.S2C,D)**. The lipid droplet protein, perilipin 2 (PLIN2) co-localised with GFP-TMEM263 at the membrane surrounding the LipidTOX-positive core **(Fig.2B-D)**. The TMEM263-positive lipid droplets are larger in size compared to controls and, particularly in OA-fed cells, we observed that GFP-TMEM263 induced a striking and significant shift in the cytoplasmic distribution of lipid droplets. Specifically, over 50% cells showed a distinctive clustered, or ‘bunch-of-grapes’ phenotype compared to untransfected controls **(Fig.2B-C,F)**. Localisation of additional lipid droplet proteins, LPCAT1 and LPCAT2 to TMEM263-positive lipid droplets further confirms TMEM263 is recruited to lipid droplet membranes **(Fig.S2E)**. A similar lipid droplet accumulation and clustering was observed upon overexpression of GFP-TMEM263 in mouse embryonic fibroblasts (MEFs) and retinal epithelial cells (RPE-1) in both BSA and OA-fed conditions **(Fig.2G,H, Fig.S2F-I)**. To confirm TMEM263 localisation to lipid droplets is not an artifact of overexpression, we observed endogenous TMEM263 localises to discrete puncta, typically 1-3 puncta per lipid droplet, at the periphery of LipidTOX-positive lipid droplets **(Fig.2I,J)**. Based on these data, we conclude that TMEM263 is an ER-resident protein which transitions to lipid droplets and the overexpression of TMEM263 is sufficient to promote the accumulation of enlarged lipid droplets with a propensity to cluster. These data are consistent with the identification of TMEM263 as a high confidence, but an uncharacterised lipid droplet protein in the literature^12, 33^.

Based on the observed subcellular localisation, we hypothesised that TMEM263 may be a class I lipid droplet protein, similar to PLIN1 and UBXD8/FAF2 which transition to lipid droplet monolayers from the ER via the ERTOLD pathway^10, 34^. These can be differentiated from proteins such as PLIN2 and ATGL1, which are recruited to lipid droplets from the cytoplasm (via cytoplasm-to-lipid droplet (CYTOLD) pathway) and are known as class II lipid droplet proteins^10, 34^. Class I proteins are classically monotopic integral membrane proteins in the ER phospholipid bilayer and have cytoplasmic N- and C-terminal tails^34, 35^. Class I proteins often insert in the ER membrane as hydrophobic hairpins, exemplified by UBXD8^36^, or alternatively via unconventional integral membrane segments, e.g. in the case of PLIN1^19^. Consistent with TMEM263 being a class I protein, it is predicted to contain 2 transmembrane domains (TM), formed by amino acid residues 38-60 and 80-102, which are well conserved across vertebrate species (**Fig.3A and Fig.S3A** and ^23, 24^). Overexpression of protein truncations of GFP-TMEM263 revealed that both TM domains are required for proper localisation of TMEM263 to the ER and to induce the characteristic accumulation of TMEM263-positive lipid droplets **(Fig.3B-D)**. Instead, constructs encoding either TM domain alone (i.e. amino acids 1-80 (TM1) and 60-116 (TM2)) showed a diffuse cytoplasmic distribution **(Fig.3C)**. Consistently, the TMEM263 mutation associated with dwarfism in chickens, W59*, which introduces an early stop codon and therefore results in the loss of TM2, shows a cytoplasmic distribution and fails to localise to the ER or lipid droplets **(Fig.3E,F)**. Deletion of the C-terminal tail, TMEM263^1-108^ (which contains the COPI-binding motif), led to a significant reduction in % cells with TMEM263-positive lipid droplets (from ∼90% to ∼45%) supporting an important functional role for the cytoplasmic C-terminus of TMEM263. Deletion of the cytoplasmic N-terminus, TMEM263^24-116^ (human isoform b) and TMEM263^33-116^, however had no impact on TMEM263 localisation to the ER, its ability to induce TMEM263-positive lipid droplets or to promote clustering of lipid droplets **(Fig.3B-D and Fig.S3B-D)**.

Despite the predicted hairpin structure, the TMEM263 AlphaFold2 (AF2) model (AF-Q8WUH6-F1) hosted on the EBI database (https://alphafold.ebi.ac.uk/entry/Q8WUH6) suggests that it exists an almost entirely unstructured conformation. To probe this further, we used ColabFold to generate 40 AF2 predictions of TMEM263 with the first 33 residues removed, since we found this region to be functionally redundant. Our data reveal that the top 5 ranked models are structurally similar, with TMEM263 folding, as predicted into a TM hairpin structure **(Fig. 3G)**. Elegant work has characterised several key biochemical determinates that support class I hairpin protein localisation to ER and transition to lipid droplets^35^. Consistent with these data, we observed several positively charged lysine residues in the TMEM263 hairpin, including at the base of the TM domains (K63, K69, K84) **(Fig.S3A,E)**, which may anchor the protein in the ER bilayer by interacting with the phospholipid head groups. This in turn forces bulky aromatic residues, in the case of TMEM263-F43 and W59 into the hydrophobic core of the bilayer **(Fig.S3A,F)**^35^. Together with the immunofluorescence observations that TMEM263 can localise to both ER and lipid droplets (Fig.2), and the published literature that TMEM263 is present in isolated lipid droplet proteomic datasets^12, 33^, these data indicate that TMEM263 may be a class I lipid droplet protein.

Given that TMEM263 is an ER-resident protein **(Fig.1C)** and its overexpression is sufficient to drive lipid droplet accumulation **(Fig.2B-E)**, we reasoned that it may be acting at the very earliest stages of lipid droplet formation, specifically lipid ‘lens’ formation in the ER bilayer. Evidence of co-localisation of endogenous TMEM263 with seipin/BSCL2, a key lipid droplet biogenesis regulator^10^, on a small proportion of lipid droplets provides support for this hypothesis **(Fig.4A)**. To further explore how TMEM263 may function at these early stages of lipid droplet biogenesis, we used coarse-grained simulations to model lens formation by introducing neutral lipids, 6% each of TAG, DAG and cholesterol esters in a bilayer. At these high concentrations, the neutral lipids naturally coalesced from their original sites, laid down in a 3×3 grid, into a ‘lens’ over the course of the 7μs simulation **(Fig.S4A**, top row). To investigate how TMEM263 behaved during lens formation, we introduced nine copies of TMEM263 (i.e. one copy per region of the 3×3 grid) into the simulation. Specifically, the top ranked AF2 model of TMEM263^33-116^ **(Fig.3G)**, which we established is indistinguishable from full-length TMEM263 in cell-based assays (**Fig.3B-D and Fig.S3B-D)** was used in all simulations. In all repeats, TMEM263 was consistently recruited to the lens border, in the same orientation, where it remained throughout the simulation. Furthermore, TMEM263 remained evenly spaced around the lens border and there was no evidence of protein-protein interactions or protein oligomerisation **(Fig.S4A)**. Consistently, immunoprecipitation with differentially tagged TMEM263 proteins showed no evidence of TMEM263 being able to bind to itself **(Fig.S4B)**.

**Figure 2.**
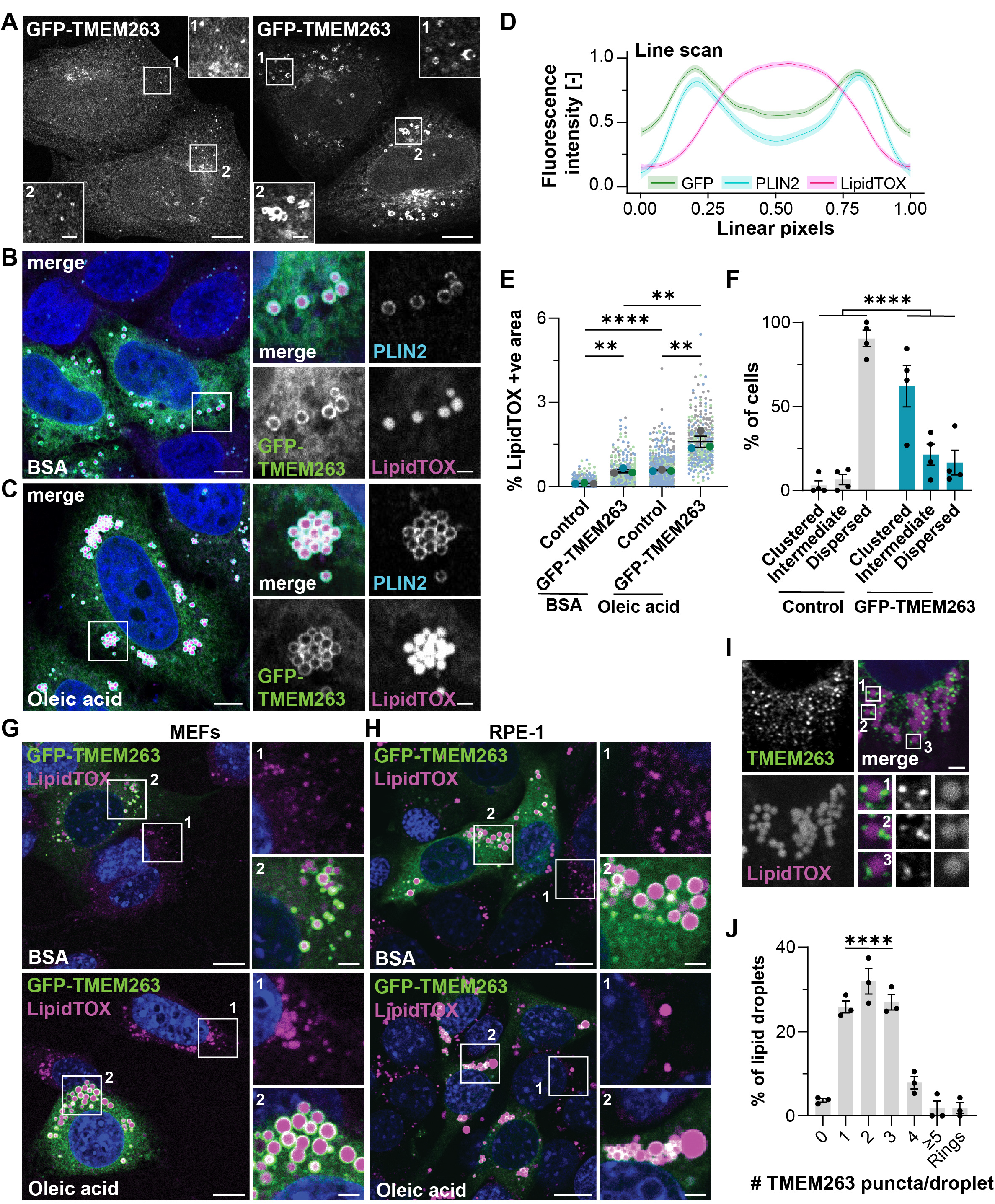
TMEM263 is a lipid droplet protein. **(A)** TMEM263 expression induces heterogenous vesicle formation. Representation images of HeLa cells expressing GFP-TMEM263. **(B,C)** TMEM263 expression induces the accumulation of lipid droplets. HeLa cells expressing GFP-TMEM263 were incubated for 16 hours with BSA vehicle control **(B)** or 30µM oleic acid **(C)**, fixed and stained with the neutral lipid dye, LipidTOX and immunostained with an antibody against endogenous perilipin 2 (PLIN2). **(D)** TMEM263 colocalises with PLIN2 at lipid droplet membranes. Quantification of GFP, PLIN2 and LipidTOX signal from 30 linescans through individual puncta. Shading corresponds to 95% confidence interval, data fitted using Loess regression. **(E)** TMEM263 induces the accumulation of lipid droplets. The % LipidTOX positive area per cell was quantified from HeLa cells treated as in B-C. Control represents untransfected cells in the same field of view as GFP-TMEM263-expressing cell. Representative images in Fig.S2 C,D. At least 250 cells per condition across n=3. Two-tailed unpaired student t-tests were performed. **(F)** TMEM263 induces clustering of lipid droplets. Cells treated as in B,C were scored as having a clustered, intermediate, or dispersed distribution of GFP-TMEM263-positive lipid droplets. Each category is expressed as a percentage of total cells. n=4, at least 150 cells per condition. Two-sided chi-squared test was performed. **(G,H)** TMEM263 induces the accumulation and clustering of lipid droplets in multiple cell lines. Mouse Embryonic Fibroblasts (MEFs) **(G)** and Retinal Epithelial (RPE-1)**(H)** cells were treated as in B,C, fixed and stained with LipidTOX. **(I,J)** Endogenous TMEM263 localises to lipid droplets. HeLa cells were treated with 30µM oleic acid for 16 hours, were fixed and immunostained with antibody against endogenous TMEM263 and stained with LipidTOX **(J)**. The number of TMEM263-positive puncta per lipid droplet was scored and is presented as a of total lipid droplets **(I)**. At least 200 lipid droplets per repeat. Ordinary one-way ANOVA with Dunnett’s multiple comparisons test was performed. Unless stated otherwise, n = 3. Scale bars are 10 µm and 2 μm for the zoomed in images. For I, scale bar is 2 μm. Unless stated otherwise, bars extend to the mean and error bars represent SEM. For E, colour coded individual data points correspond to individual cells from different experiments, lines show mean values, and error bars represent SEM. ns p ≥ 0.05, * p < 0.05,** p < 0.01, *** p < 0.001.

These data are consistent with previous reports, that hairpin proteins have affinity for TAG and can localise around a neutral lipid lens in a bilayer^37^. Coupled with the fact that all copies of TMEM263 were in the same orientation towards the lipids, this led us to hypothesise that specific residues in TMEM263 may bind directly to lipids. To provide a higher resolution view of this, we carried out atomistic simulations of TMEM263 in a bilayer containing TAG and DAG, and used the PyLipID package to assess the neutral lipid binding properties of individual residues^38^. This analysis shows there is a clear preference for TAG and DAG binding to the transmembrane domains of TMEM263 **(Fig.4B and Fig.S4C)**. Interestingly, the hydrophobic TM domains of seipin/BSCL2 have been shown in molecular dynamics simulations to trap and cluster TAG and DAG in a bilayer, concentrating the lipids and lowering the energy barrier to lens formation^16^. To test whether TMEM263-lipid binding has a similar role and is functionally important in promoting lipid lens formation, coarse-grained simulations were repeated using a lower concentration of 2.5% TAG in the bilayer, and no other neutral lipids **(Fig. 4C)**. This concentration is below the threshold at which lipids would coalesce spontaneously, as seen from our no protein control **(Fig. 4C**, left column) and thus allowed us to test whether TMEM263 could contribute to lens formation. The data showed that one molecule of TMEM263 was sufficient to induce a significant increase in the proportion of TAG within the lens, supporting the hypothesis that TMEM263 can promote lipid droplet biogenesis **(Fig. 4C,D)**.

The first step to experimentally validate this hypothesis, was to further interrogate the atomistic simulations and PyLipID analysis **(Fig.4B and Fig.S4C)** to identify the specific amino acids important for lipid binding. We observed that both neutral lipids, TAG and DAG accumulate within the centre of TMEM263 during the atomistic simulations (**Fig.S4D)**, making especially high contact to residue W59 and K84, and more weakly, F43, suggesting that these residues may facilitate lipid binding (**Fig.4E,F and Fig.S4C**). The three putative lipid-binding residues were mutated to alanine and overexpressed in HeLa cells. In all cases, the percentage of cells expressing GFP-TMEM263 positive puncta was significantly reduced compared to wild-type control **(Fig.4G)**. Co-staining with the ER marker, mCherry-Sec61b confirmed that F34A and K84A were still correctly localised to ER, while W59A showed a cytoplasmic distribution **(Fig.S4E)**. As it is not localised properly, it would not be possible to interrogate the importance of W59 lipid-binding, independent of ER localisation and therefore the W59A mutant was omitted from subsequent analysis. Wild-type TMEM263 induced a significant increase in the LipidTOX-positive area compared to transfected controls in both BSA control and OA-fed conditions, however F34A and K84A failed to promote an increase in the LipidTOX-positive area above the untransfected controls (**Fig.4H and Fig.S4F**). Furthermore, neither F43A nor K84A induced a change in the distribution of lipid droplets from dispersed to clustered as seen upon expression of wild-type TMEM263 (**Fig.S4F,G**).

Together, these data suggests that TMEM263 can bind neutral lipids in the ER bilayer, potentially to aid lens formation. Our data indicates that TMEM263 can then transition onto the monolayer of the growing lipid droplets. These data would suggest TMEM263 is required for efficient lipid droplet biogenesis, and we therefore hypothesised that loss or mutation of TMEM263 would conversely impair lipid droplet formation. To interrogate this hypothesis, we generated CRISPR/Cas9 knock-out (KO) HeLa cell lines **(Fig.S5A)** and CRISPR/Cas9 mutant zebrafish lines **(Fig.S5B,C)**. Two clonal *Tmem263* KO HeLa lines showed significantly lower BODIPY-positive area **(Fig.5A,B)** and significantly reduced lipid droplet numbers **(Fig.5A,C)** compared to parental controls following OA feeding. Interestingly, the majority of *Tmem263* crispant fish (77.5%) showed abnormal phenotypes, including oedema, bent tails and tail truncations **(Fig.S5D,E)**, and failed to survive past 5 days post fertilisation (dpf). The surviving 22.5% *Tmem263* crispant fish showed a small but significant decrease in body length **(Fig.5D,E)**, consistent with an established role for TMEM263 in organismal growth^23-25^. Lipid droplet accumulation in the intestine was significantly decreased in TMEM263 crispants following OA feeding **(Fig.5F,G)**. It is important to note that given that experiments could only be carried out in 22.5% of *tmem263* crispants with less severe phenotypes, it is likely the results presented here are under-representing the extent of the growth and lipi droplet phenotype. Altogether, these data demonstrate TMEM263 is necessary for efficient lipid droplet formation *in vivo*. These results provide a novel and previously unrecognised link between lipid droplet homeostasis and organismal growth.

## Discussion

Using *in silico, in cellulo* and *in vivo* models, we have identified TMEM263 as an ER-resident protein, and a COPI cargo which is necessary and sufficient for efficient lipid droplet formation. Our data demonstrate that TMEM263 overexpression causes the accumulation of enlarged lipid droplets that have a propensity to cluster, particularly upon lipid feeding, and are enriched with classical lipid droplet proteins including perilipin 2 and LPCATs. Molecular dynamics simulations revealed that TMEM263 can bind to and concentrate neutral lipids in a phospholipid bilayer. Mutation of these residues impairs TMEM263-induced lipid droplet formation. Based on these data we propose that TMEM263 lowers the energy barrier to lipid lens formation in the ER and contributes to lipid droplet formation. Consistently, loss or mutation of TMEM263 in cells and *in vivo*, impairs lipid droplet formation.

Data presented here suggests that TMEM263 has many biochemical features of a class I lipid droplet protein – including the ability to transition from the ER phospholipid bilayer to the lipid droplet monolayer. The latter localisation is further supported in the literature by the identification of TMEM263 in proximity labelling of the lipid droplet surface^12^, and in isolated lipid droplets treated with ionic chaotropes (to specifically preserve monolayer integrated proteins as opposed to peripherally associated proteins)^33^. Interestingly, the latter study, TMEM263 (or C12orf23) was not identified to contain a predicted hydrophobic membrane domain, or a predicted lipid-anchoring motif and it is unclear from our modelling whether the TM domains span the entire bilayer or may be a monotopic hairpin thus it is currently unclear how TMEM263 integrates into the monolayer. One possibility is that TMEM263 may behave similarly to UBXD8/FAF2, one of the best characterised class I lipid droplet proteins, which reorganises from a deeply inserted V-shaped topology in the ER bilayer (similar to TMEM263 in our coarse-grained simulations) into a shallower hairpin in the monolayer^36^. However, the helices in UBXD8 are substantially shorter than seen in our TMEM263 model (ca. 12 residues vs ca. 22). Alternatively, it may transition out of the bilayer to associate with the monolayer via an amphipathic helix, or transition may be facilitated via an unknown binding partner. We found no evidence that TMEM263 can form any higher order oligomers, but that does not preclude TMEM263 has other functionally important binding partners. Indeed, given that TMEM263 is such a small protein, with no obvious functional domains, it may act either in the ER or on lipid droplets as a scaffold, or an anchoring protein. It will be important to explore if TMEM263 has location-specific (i.e. ER vs lipid droplet) binding partners and functional roles. For example, given our hypothesis that TMEM263 functions to concentrate lipids in the ER bilayer, it remains to be seen what the function of TMEM263 may be on the lipid droplet monolayer. The distinctive clustering of lipid droplets upon TMEM263 overexpression may hint at a functional role for the protein in regulating membrane contact sites between lipid droplets or between lipid droplets and the ER.

Similar to TMEM263, the hydrophobic transmembrane domains of the multimeric lipid droplet biogenesis regulator, seipin/BSCL2 can also cluster TAG and DAG in molecular dynamics simulations ^16^, potentially indicating there may be mechanistic commonalities between the proteins. Interestingly, there is limited evidence of co-localisation between seipin and TMEM263 in our experiments, and no evidence of interaction by immunoprecipitation (data not shown) so it remains an interesting open question how TMEM263 coordinates with the established mechanisms to control the very earliest stages of lipid droplet formation. One possibility is that TMEM263 may be involved in the dynamic contacts that lipid droplets maintain with the ER to facilitate the transfer of lipids (and proteins) in response to changes in metabolic demand. Potential support for this hypothesis is that unlike seipin, where usually only one punctum is observed per lipid droplet (i.e. the site of biogenesis)^39^, we frequently see 2 or 3 TMEM263-positive puncta per lipid droplet. Lipid droplets maintain a dynamic physical and functional association with ER^20^, and our data may indicate TMEM263 is involved in more extensive ER-lipid droplet contact sites, possibly controlling the delivery of lipids in a pathway parallel to seipin. Alternatively, TMEM263 may be involved in controlling lipid droplet size rather than biogenesis *per se*. The late emergence of TMEM263, which is limited to vertebrates, is in contrast to seipin which is evolutionarily conserved^14^ and may reflect the increased regulatory complexity required in vertebrates to control lipid storage and metabolism.

While there are still questions about the role and molecular regulation of TMEM263, the biggest mystery is how loss or mutation of TMEM263, an ER-resident protein and lipid droplet regulator, causes a severe impairment of organismal growth and bone health. Human GWAS studies have shown a significant association between TMEM263 variants and bone mineral density, bone fracture risk and osteoblast function^21, 22^. Introduction of early stop codons in TMEM263 causes autosomal dwarfism in chickens^24^ and was linked to a case of severe fatal skeletal dysplasia and shortened limbs in a human foetus^25^. Data presented in our study suggests that in both cases, these stop codons would result in loss of TMEM263 TM2 domain and functionally inactivate the protein. Consistently, TMEM263 KO mice showed proportional dwarfism, and were 30-40% smaller than wild-type littermates at 8 weeks old^23^. While the physiological consequences of TMEM263 loss are consistent across model species, there is no clear causative mechanism. In contrast to the observations in humans and chickens for example, the KO mice were born at the same weight and size as wildtypes, with growth defects appearing after 21 days, a period of intense long bone growth^23^. Furthermore, while the TMEM263 mutant chickens show no changes in plasma IGF1 levels^31, 32^, the TMEM263 KO mice showed reduced circulating IGF1 levels, possibly a result of reduced GHR expression and reduced JAK2/STAT5 signalling in the liver^23^. There was no mention of lipid defects or lipodystrophy in any of these reports, however transcriptomics from the TMEM263 KO mouse livers identified fatty acid metabolism and steroid hydroxylase activity were among the most differentially expressed genes (GO terms), indicating that lipid handling and metabolism is grossly altered in these animals^23^. More extensive analysis is required of the model organisms, the KO mice and the zebrafish models we have generated here to explore lipid homeostasis and determine if there are changes in physiological resilience in response to lipid or metabolic challenges. Our data identified a clear cell-autonomous role for TMEM263 in the regulation of lipid droplets and evidence in mouse and human tissue suggests TMEM263 is ubiquitously expressed in all tissues^23^. It therefore remains to be seen whether the whole-body defects in growth are a result of cell or tissue-specific roles.

The ability to sense and properly respond to changes in nutrient availability is fundamental to regulate growth. While we have a good understanding of how growth factors and amino acids integrate to promote growth pathways, such as mTORC1 and the autophagy-lysosome pathway, much less is known about the links between lipid homeostasis and growth. Dysregulation of lipid droplets and ER-lipid droplet crosstalk is widely linked to metabolic diseases such as obesity, non-alcoholic fatty liver disease (e.g., PNPLA3, PLIN3), and lipodystrophy (e.g., seipin/BSCL2, AGPAT2)^11, 40^. This is however one of the first reports of lipid droplets being linked to the regulation whole-body growth and bone health. Previous studies have identified lytic bone lesions in patients with AGPAT mutations ^41^; short stature^42^ and possibly GH deficiency^43^ have been noted in patients with mutations in the lipophagy regulator, spartin^42^, and mutations in CAV1 or PTRF genes, which result in congenital generalised lipodystrophy, have been associated with short stature and skeletal abnormalities ^44 45, 46^. In all cases however the mechanisms are poorly understood. Thus, in TMEM263, we have identified a new fundamental mechanistic link between lipid homeostasis and organismal growth.

## Acknowledgements

BC lab (BC, KY, NC, PZ, LW, SW and TA) is supported by The Royal Society and Wellcome Trust (218547/Z/19/Z), BBSRC (BB/Y001427/1) and AMS Springboard Award (SBF005\1130). JM and CH are supported by BBSRC (BB/Y002504/1). The authors gratefully acknowledge the Wolfson Bioimaging Facility, the Bristol Proteomics Facility, and the Animal Service Unit (ASU), University of Bristol, UK, for their support and assistance in this work. The authors thank Rachel Curnock and Ben Clark for their technical assistance with cloning. This work was carried out using the computational facilities of the Advanced Computing Research Centre, University of Bristol -http://www.bristol.ac.uk/acrc/. This project made use of time on ARCHER2 granted via the UK High-End Computing Consortium for Biomolecular Simulation, HECBioSim (http://hecbiosim.ac.uk), supported by EPSRC (EP/R029407/1).

## Figure Legends

**Figure S1-Identification of COPI cargo proteins. (A,B)** GFP-εCOPI interacts and localises with endogenous COPI complexes. HeLa cells stably expressing GFP-εCOPI were subject to GFP immunoprecipitation and western blot to confirm binding with endogenous COPI complexes **(A)** and colocalises with endogenous Sec61β (ER) and TGN46 (trans-Golgi) **(B)**. n = 2. Scale bars: 10 μm. **(C)** Identification of COPI cargo involved in growth. HeLa cells stably expressing GFP or GFP-εCOPI were subject to GFP immunoprecipitation and analysis by TMT mass spectrometry. Volcano plot of proteins enriched in GFP-ε-COPI / GFP immunoprecipitation, proteins in teal represent COPI cargoes, defined as proteins with a log fold change >2.5, p-value <0.005 GFP-εCOPI / GFP (n = 4). Proteins labelled on the volcano plot are the top 10 proteins with canonical COPI-interacting motifs, and those in pink are specifically associated with mammalian growth regulation. **(D)** Validation of GFP-εCOPI interaction with TMEM263. HeLa cells overexpressing mCherry-TMEM263 and GFP-ε-COPI were subject to GFP immunoprecipitation and western blotting. n = 2.

**Figure S2-Characterising TMEM263 positive vesicles. (A)** TMEM263 expression induces puncta formation. Quantification of the % HeLa cells expressing GFP or GFP-TMEM263 that contain GFP puncta. At least 600 cells per condition across n=3. Two-sided chi-squared test was performed. **(B)** GFP-TMEM263 puncta are lipid droplets. Quantification of the % HeLa cells with LipidTOX-positive GFP-TMEM263 puncta. At least 100 cells, at least 1500 puncta per condition across n=3. **(C, D)** Expression of TMEM263 induces the accumulation and clustering of lipid droplets. HeLa cells expressing GFP-TMEM263 were incubated for 16 hours with BSA vehicle control **(C)** or 30µM oleic acid **(D)**, fixed and stained with LipidTOX. White dotted line highlights GFP-transfected cells which were quantified relative to neighbouring untransfected controls in the same field of view. Quantification in Fig.2E. **(E)** TMEM263 colocalises with lipid droplet proteins. HeLa cells expressing GFP-TMEM263 were fixed and immunostained with antibodies against endogenous LPCAT1 and LPCAT2. **(F-I)** TMEM263 induces the accumulation and clustering of lipid droplets in multiple cell lines. Mouse Embryonic Fibroblasts (MEFs) **(F,G)** and Retinal Epithelial (RPE-1) **(H,I)** cells were treated as C-D, fixed and stained with LipidTOX. The % LipidTOX positive area per cell and the % cells with clustered, intermediate or dispersed GFP-TMEM263-positive lipid droplet distribution were quantified. For F, 2 outliers are out of range of axis.

Unless stated otherwise, n = 3. Scale bars are 10 µm and 2 μm for the zoomed in images. Bars extend to the mean and error bars represent SEM. For F and H, colour coded individual data points correspond to individual cells from different experiments, lines show mean values, and error bars represent SEM. ns p ≥ 0.05, * p < 0.05, ** p < 0.01, *** p < 0.001, **** p < 0.0001.

**Figure S3-Identifying functional domains of TMEM263. (A)** Multiple sequence alignment of TMEM263 highlights the two transmembrane domains are highly conserved. **(B-D)** N-terminus of TMEM263 is functionally redundant. HeLa cells expressing GFP-TMEM263 full-length or GFP-TMEM263^33-116^ were incubated for 16 hours with BSA vehicle control or 30µM oleic acid, fixed and stained with the neutral lipid dye, LipidTOX **(B)**. The % LipidTOX positive area per cell was quantified **(C)**. Ordinary one-way ANOVA with Dunnett’s multiple comparisons tests and two-tailed unpaired student t-tests were performed. Control represents untransfected cells in the same field of view as GFP-TMEM263-expressing cells. Data from control and GFP-TMEM263^WT^-expressing cells are also included in Fig. 2E. Cells were scored as having a clustered, intermediate, or dispersed distribution of GFP-TMEM263-positive lipid droplets. Each category is expressed as a percentage of total cells **(D)**. At least 200 cells per condition across n=3. Two-sided chi-squared tests were performed. **(E-F)** TMEM263 shares biochemical features with class I lipid droplet proteins. TMEM263^33-116^ AlphaFold2 model (magenta) modelled in a phospholipid bilayer with lysines labelled in purple **(E) and** aromatic residues labelled in blue **(F)**. Unless stated otherwise, n=3. Scale bars are 10 µm and 2 μm for the zoomed in images, bars extend to the mean, colour coded individual data points correspond to individual cells from different experiments, lines show mean values, and error bars represent SEM. ns p ≥ 0.05, * p < 0.05, ** p < 0.01, *** p < 0.001, **** p < 0.0001.

**Figure S4-Characterising TMEM263 lipid binding. A)** TMEM263 is recruited to lipid lens in a phospholipid bilayer. Coarse-grained simulation of 6% TAG (green), 6% DAG (cyan), and 6% cholesterol esters (blue) in a phospholipid bilayer with 9 molecules of TMEM263^33-116^ (magenta). Phosphate heads are shown in grey in the side view. Views are shown both before (0µs) and after (7µs) simulation. **(B)** TMEM263 shows no evidence of oligomerisation. HEK293T cells expressing GFP-TMEM263 and HA-TMEM263 were subject to GFP immunoprecipitation and western blot. **(C)** Binding frequencies for each residue in TMEM263 for TAG and DAG. These are the same data as shown in Fig. 4B. **(D)** Snapshots showing neutral lipids moving around the TMEM263 model based on atomistic simulation data. **(E)** Lipid binding mutants do not impact TMEM263 localisation to the ER. HeLa cells stably expressing mCherry-Sec61b were transfected with GFP-TMEM263 mutants as indicated. **(F,G)** HeLa cells were transfected with GFP-TMEM263 mutants as indicated, were incubated for 16 hours with BSA vehicle control or 30µM oleic acid, fixed and stained with the neutral lipid dye, LipidTOX. Cells were scored as having clustered, intermediate, or dispersed lipid droplets. At least 250 cells per condition across n=3. Two-sided chi-squared tests were performed. Data from control and GFP-TMEM263^WT^-expressing cells were also included in Fig. S3D.

Unless stated otherwise, n=3. Scale bars are 10 µm and 2 μm for the zoomed in images, bars extend to the mean and error bars represent SEM. * p < 0.05, **** p < 0.0001.

**Figure S5-Generation of TMEM263 models. (A)** Generation of TMEM263 Crispr/Cas9 knock-out (KO) HeLa cells. Representative blot showing loss of TMEM263 protein. **(B,C)** Strategy to create TMEM263 mutant zebrafish. gRNA were generated to target the transmembrane domains of TMEM263 **(B)** and the timeline to create a crispant zebrafish cohort **(C). (D,E)** TMEM263 mutation impairs zebrafish larval development. The proportion of zebrafish in control or TMEM263 crispant groups were quantified for normal or abnormal larval development **(D)**. Representative image of two TMEM263 crispant larvae (5dpf) with characteristic oedema (black arrow) **(E)**.

## Materials and Methods

### Cell lines and treatments

Human cervical cancer HeLa cells, HEK293T cells, wild-type mouse embryonic fibroblasts (MEF), and RPE1-hTERT cells were cultured in DMEM (D6546, Sigma-Aldrich) supplemented with 2 mM glutamine, 10% FBS and 100 U/ml penicillin–streptomycin at 37°C, 5% CO_2_. Where indicated, 24 hours post-transfection, cells were incubated for 16 hours in complete DMEM supplemented with 30µM oleic acid (O1383, Sigma-Aldrich) in 10% fatty acid-free BSA (A8806, Sigma-Aldrich) or 10% fatty acid free BSA only as vehicle control.

### Generation of TMEM263 KO cell lines

gRNA targeting TMEM263 was cloned into pSpCas9(BB)-2A-Puro using Bbs1-HF. gRNA 1: GGUAUCUUCAGUGUUACAAA (to generate TMEM263 KO1) and 2: CUUAAUGAUGAACCACCAGA (to generate TMEM263 KO2). HeLa cells were transfected with FuGENE HD (Promega) overnight and selected with 3ug/ml puromycin for 48 hours. To generate single clones, cells were subsequently subject to serial dilution. Clones were screened by western blot.

### DNA and viral production

mCherry-Sec61b (addgene #49155) was subcloned into pXLG3 (SpeI and KpnI). pLVXpuro-εCOPI-GFP was a gift from Prof David Stephens, University of Bristol (addgene #66604). TMEM263 cDNA (NP_001306590.1) was purchased from Integrated DNA Technologies (IDT) and cloned into pEGFP-C2 (Clontech), p-mCherry2-C1 (addgene #54563)), FLAG-HA-pcDNA3.1 (addgene #52535). Primers used to generate TMEM263 truncations and point mutations in pEGFP-C2, are shown in **Table 1**. Point mutants were generated using QuikChange XL site-directed mutagenesis kit (Agilent 200517). All transient transfections were performed using FuGENE6 transfection reagent (Promega E2691) or Lipofectamine2000 according to manufacturer instructions. Transient transfection of HEK293T cells was carried out using 10 μg/μl polyethylenimine (PEI) with a 3:1 ratio PEI:DNA.

For virus production, HEK293T cells were transfected with plasmid containing the gene of interest and packaging vectors pAX2 and pMDG2 for 72 hours using 10 μg/μl PEI with a 3:1 ratio PEI:DNA. Viruses were harvested, filtered and incubated on cells overnight.

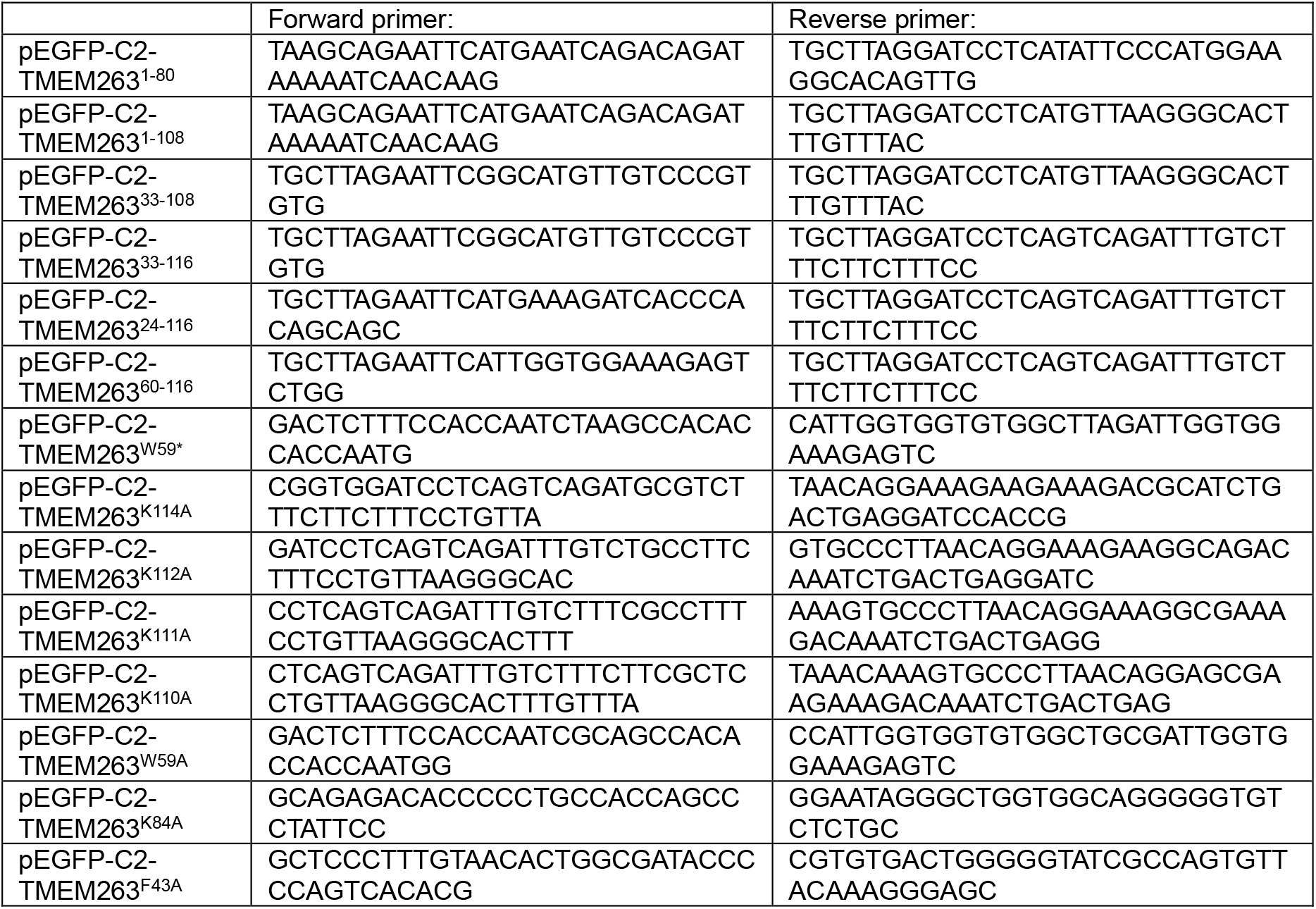

### Zebrafish husbandry and CRISPR-mutant lines

Zebrafish (*Danio rerio*) were raised and maintained under standard conditions^47^. Experiments were approved by the local ethics committee (the Animal Welfare and Ethical Review Body of the University of Bristol) and performed under UK Home Office project licence.

Crispant *tmem263* zebrafish lines were generated using CRISPR-Cas9 mutagenesis, as previously performed^48^. Briefly, gRNAs targeting exon 3 of *tmem263* (ENSDARG00000030465) were synthesised (2nmol, Alt-R™ CRISPR Cas9 crRNA, IDT): CRISPR-guide 1 – ATGGGAGTAGGCCTGGTGAAGGG; CRISPR-guide 2 – GACCATCCCCAGCCTCAGCCTGG, CRISPR-guide 3 –GGCGCTGTTGGGGCTACAGTGGG. crRNAs were each annealed to Alt-R CRISPR-Cas9 tracrRNA (5nmol, IDT) at 95°C for 5 minutes to form a 57mM gRNA. Each gRNA was then incubated with Alt-RTM S.p. Cas9 Nuclease V3 (diluted to 57mM, IDT) for 5 minutes at 37°C to assemble the ribonucleoprotein (RNP) for injection. The yolk of 1-cell stage zebrafish embryos was injected with 1nl of RNP (1000pg gRNA and 4700pg of Cas9) and surviving embryos were raised under normal husbandry conditions. All three gRNAs were injected in combination.

### DNA extraction and genotyping of zebrafish lines

For DNA extraction, whole embryos were incubated in base solution (25 mM NaOH, 0.2 mM EDTA) for 30 minutes at 95°C before addition of equal volume of neutralisation solution (40 mM Tris–HCl, pH 5.0). For genotyping, DNA were amplified using the following primers: F-TGCAACCTGCATCTTATGGAGA, R-AGAGGTCCGTGTCTCTACCC. Amplicons were sent for Sanger sequencing.

### Lipid droplet analysis in zebrafish

BODIPY FL C_5_ (D3834, Invitrogen) was used for live imaging of lipid droplets in the intestines of larvae following OA treatment. Briefly, larvae were fed 300μM oleic acid in Danieau’s buffer for 24 hours from 5-6dpf, rinsed three times, then placed in 3123μM BODIPY in Danieau’s buffer for 2 hours and imaged.

### Immunofluorescence

Cells were seeded onto glass coverslips for 24-48 hours then fixed using 4% paraformaldehyde/PBS for 10 minutes and permeabilised in 0.2% TritonX-100/PBS for 8 minutes, all at room temperature. Alternatively, 0.1% saponin/PBS was as permeabilization buffer when staining for endogenous TMEM263 and seipin, and 0.01% saponin added to all subsequent incubation and wash solutions. Coverslips were incubated with blocking buffer (5% BSA/PBS-0.05% Tween) for 1 hour at room temperature. Coverslips were incubated in primary antibodies in blocking solution overnight at 4°C. Primary antibodies used in this study:γCOP (Proteintech, 12393-1-AP, rabbit, 1:1000), ADRP/Perilipin-2 (Proteintech, 15294-1-AP, rabbit 1:400), TMEM263 (Invitrogen, PA5-70960, rabbit 1:200 & NovusBio, BP2-83690, rabbit, 1:200), Sec61B (Cell Signaling Technology, #14648, rabbit, 1:200), TGN46 (GeneTex, GTX74290, sheep, 1:2000), BSCL2/Seipin (Abnova, H00026580-A02, mouse, 1:200), LPCAT1 (Abcam, ab214034, rabbit, 1:250), LPCAT2 (Abcam, ab224244, rabbit, 1:250). Coverslips were washed and incubated in the appropriate Alexa Fluor-conjugated secondary antibodies (Thermo Fisher Scientific, 1:1000) in blocking buffer for 1 hour at room temperature. The following secondary antibodies were used: goat anti-rabbit 594 (A11037, Invitrogen, Thermo Fisher Scientific), donkey anti-rabbit 568 (A10042, Invitrogen, Thermo Fisher Scientific), goat anti-rabbit 488 (A11034, Invitrogen, Thermo Fisher Scientific), donkey anti-mouse 568 (A10037, Invitrogen, Thermo Fisher Scientific), and donkey anti-sheep 633 (A21100, Invitrogen, Thermo Fisher Scientific). Where indicated, cells were incubated at room temperature with HCS LipidTOX™ Deep Red Neutral Lipid Stain (H34477, Invitrogen 1:1000) for 30 minutes or BODIPY-493/503 (25892, Cayman chemical 1µg/ml) for 10 minutes, diluted in PBS. Coverslips were mounted using ProLong™ Gold Antifade Mountant with DAPI (Invitrogen, Thermo Fisher Scientific, P36934).

### Microscopy

Cells were imaged using either a Leica SP8 AOBS confocal laser scanning microscope with 63×HC PL APO CS2 oil objective, or an Olympus IXplore SpinSR system with 60X oil objective.

For live imaging, fish were anaesthetised using 0.1 mg/ml MS222 (Tricaine methane sulfonate) diluted in Danieau’s buffer and imaged laterally. Images of live larvae at 5dpf were obtained using a Leica MZ10 F modular stereo microscope system at 3.2x magnification. Alternatively, zebrafish larvae were mounted in 1% LMP agarose (16520050, Thermofisher) and imaged using a Leica SP5-II AOBS tandem scanner confocal microscope attached to a Leica DMI 6000 inverted epifluorescence microscope and oil immersion 20x objectives run using Leica LAS AF software (Leica, Germany).

### Image quantification

The of percentage of cells containing GFP-positive puncta was scored, by a blinded researcher, manually on a widefield Leica DMI6000 inverted epifluorescence microscope. Scoring of number of TMEM263 puncta (Fig.2I), scoring of the percentage of LipidTOX positive GFP puncta (Fig.S2B), and analysis of lipid droplet distribution (i.e. dispersed, intermediate or clustered) was carried out manually post-imaging. All other image quantification was carried out using ImageJ (Fiji) software. For lipid droplet quantification (i.e. count and percentage area), a constant threshold was applied across all images, then an ROI was generated for each individual cell and the ‘analyse particles’ tool utilised with no set minimum size. For the linescan analysis of GFP/PLIN2/LipidTOX (Fig.2C), 30 lipid droplets from 1 independent experiment were analysed. A line was drawn across individual lipid droplets and using the ‘Plot Profile’ function of ImageJ/Fiji the raw fluorescent intensities of the 3 channels were determined. The fluorescent intensities were then normalised to the maximum fluorescent intensity and longest distance of each channel and lipid droplet.

For zebrafish, standard larval lengths were obtained from lateral images at 5 dpf by measuring from the nose of the fish to the end of the tail manually in Fiji. For lipid droplet analysis, a 68 μm2 region of interest (ROI) was selected from one z-slice per larvae and BODIPY puncta counted. Maximum projection images were assembled using LAS AF Lite software (Leica) and Fiji (Schindelin et al., 2012) (Fig.5F).

### Immunoprecipitation

HEK293T were seeded in 10cm dishes and lysed for immunoprecipitation (IP) 24hours after transfection. Cells were washed in ice-cold PBS and lysed, on ice, in 500μl IP buffer (40mM Tris-HCL pH 7.5, 0.5% NP-40 (IGEPAL CA-630, #18896, Sigma-Aldrich), supplemented with 1x Halt™ Protease and Phosphatase Inhibitor Cocktail, EDTA-free (Thermo Fisher Scientific, 78443,1:100). Lysates were cleared by centrifugation for 10 minutes at 13,500rpm at 4°C. A whole cell lysate sample was removed and the remaining supernatant was incubated with 10μl of equilibrated GFP-Trap agarose (GTA-20, ChromoTek) for 2 hours at 4°C with constant rotation, then washed and samples boiled at 95°C for 5 minutes in 1xNuPAGE LDS sample buffer (NP0008, Invitrogen, Thermo Fisher Scientific) supplemented with 5% 2-mercaptoethanol.

### Immunoblotting

Immunoblotting was performed as described previously^49, 50^. Briefly, samples were separated via SDS-PAGE using 4–12% NuPAGE gels, (Thermo Fisher Scientific, NP0335BOX and NP0336BOX) as per company instructions. Gels were transferred to methanol-activated PVDF membrane by either standard wet transfer or semi-dry transfer methods. Membranes were blocked for one hour at room temperature in 5% milk in 0.05% PBS-Tween. Primary antibodies were diluted in blocking buffer and incubated overnight at 4°C. The following primary antibodies were used: HA.11 (Biolegend, #901573, mouse, 1:1000), GFP (Roche, 11814460001, mouse, 1:2000), mCherry (Abcam, ab167453, rabbit, 1:2000), TMEM263 (Invitrogen, PA5-70960, rabbit 1:400), γCOP (Proteintech, # 12393-1-AP, rabbit, 1:1000),βCOP (gift from D. Stephens, University of Bristol, mouse, 1:500), and GAPDH (Cell Signaling Technology, #97166, mouse, 1:2000). After 3 washes with PBS-T, the blots were incubated for one hour at room temperature with the following secondary antibodies diluted in blocking buffer: goat anti-mouse 680 (Invitrogen, Thermo Fisher Scientific, A21057, 1:20000) and goat anti-rabbit 800 (Invitrogen, Thermo Fisher Scientific, SA535571, 1:20000). Blots were washed 3 times with PBS-T and once with PBS before being imaged with LI-COR Odyssey xL (LI-COR Biotechnology) and quantified using Image Studio Lite software (LI-COR Biotechnology).

### Proteomics

#### TMT Labelling and High pH reversed-phase chromatography

Immuno-isolated samples were reduced (10mM TCEP, 55°C for 1h), alkylated (18.75mM iodoacetamide, room temperature for 30min.) and then digested from the beads with trypsin (1.25µg trypsin; 37°C, overnight). The resulting peptides were then labeled with TMTpro sixteen-plex reagents according to the manufacturer’s protocol (Thermo Fisher Scientific, Loughborough, LE11 5RG, UK) and the labelled samples pooled and desalted using a SepPak cartridge according to the manufacturer’s instructions (Waters, Milford, Massachusetts, USA). Eluate from the SepPak cartridge was evaporated to dryness and resuspended in buffer A (20 mM ammonium hydroxide, pH 10) prior to fractionation by high pH reversed-phase chromatography using an Ultimate 3000 liquid chromatography system (Thermo Scientific). In brief, the sample was loaded onto an XBridge BEH C18 Column (130Å, 3.5 µm, 2.1 mm X 150 mm, Waters, UK) in buffer A and peptides eluted with an increasing gradient of buffer B (20 mM Ammonium Hydroxide in acetonitrile, pH 10) from 0-95% over 60 minutes. The resulting fractions (concatenated into 6 in total) were evaporated to dryness and resuspended in 1% formic acid prior to analysis by nano-LC MSMS using an Orbitrap Fusion Tribrid mass spectrometer (Thermo Scientific).

#### Nano-LC Mass Spectrometry

High pH RP fractions were further fractionated using an Ultimate 3000 nano-LC system in line with an Orbitrap Fusion Tribrid mass spectrometer (Thermo Scientific). In brief, peptides in 1% (vol/vol) formic acid were injected onto an Acclaim PepMap C18 nano-trap column (Thermo Scientific). After washing with 0.5% (vol/vol) acetonitrile 0.1% (vol/vol) formic acid, peptides were resolved on a 250 mm × 75 μm Acclaim PepMap C18 reverse phase analytical column (Thermo Scientific) over a 150 min organic gradient with a flow rate of 300 nl min^™1^. Solvent A was 0.1% formic acid and Solvent B was aqueous 80% acetonitrile in 0.1% formic acid. Peptides were ionized by nano-electrospray ionization at 2.0kV using a stainless-steel emitter with an internal diameter of 30 μm (Thermo Scientific) and a capillary temperature of 275°C.

All spectra were acquired using an Orbitrap Fusion Tribrid mass spectrometer controlled by Xcalibur 2.1 software (Thermo Scientific) and operated in data-dependent acquisition mode using an SPS-MS3 workflow. FTMS1 spectra were collected at a resolution of 120 000, with an automatic gain control (AGC) target of 200 000 and a max injection time of 50ms. Precursors were filtered with an intensity threshold of 5000, according to charge state (to include charge states 2-7) and with monoisotopic peak determination set to peptide. Previously interrogated precursors were excluded using a dynamic window (60s +/-10ppm). The MS2 precursors were isolated with a quadrupole isolation window of 1.2m/z. ITMS2 spectra were collected with an AGC target of 10 000, max injection time of 70ms and CID collision energy of 35%.

For FTMS3 analysis, the Orbitrap was operated at 50 000 resolution with an AGC target of 50 000 and a max injection time of 105ms. Precursors were fragmented by high energy collision dissociation (HCD) at a normalised collision energy of 60% to ensure maximal TMT reporter ion yield. Synchronous Precursor Selection (SPS) was enabled to include up to 10 MS2 fragment ions in the FTMS3 scan.

### Data Analysis

The raw data files were processed and quantified using Proteome Discoverer software v2.4 (Thermo Scientific) and searched against the UniProt Human database (downloaded January 2024: 82415 entries) using the SEQUEST HT algorithm. Peptide precursor mass tolerance was set at 10ppm, and MS/MS tolerance was set at 0.6Da. Search criteria included oxidation of methionine (+15.995Da), acetylation of the protein N-terminus (+42.011Da) and methionine loss plus acetylation of the protein N-terminus (−89.03Da) as variable modifications and carbamidomethylation of cysteine (+57.021Da) and the addition of the TMT mass tag (+304.207Da) to peptide N-termini and lysine as fixed modifications. Searches were performed with full tryptic digestion and a maximum of 2 missed cleavages were allowed. The reverse database search option was enabled and all data was filtered to satisfy false discovery rate (FDR) of 5%. COPI cargo were defined as proteins with log fold change (FC) > 2.5 and p-value <0.005 GFP-εCOPI / GFP (n = 4). Proteins were further filtered for those containing COPI-interacting motifs, either KxKxx or KKxx (at the C-terminus) or RxR (within the last 30 amino acids of C-terminus), and associated with ‘growth’ in the MGI database^27, 28^.

### Statistical analysis

For the statistical analysis of quantitative values, two kinds of parametric tests were performed on experimental data from at least three individual experiments. The two-tailed unpaired student t-test or one-way ordinary ANOVA for multiple comparison test with Dunnett’s correction was performed for comparisons between two or more than two conditions, respectively. For the statistical analysis of qualitative data (clustering, GFP puncta positive), the chi-squared two tailed test or the Fisher’s exact test was performed on individual experiments. The same test was also performed on pooled experiments and shown in graphs. All statistical tests were performed with GraphPad Prism v10.2.3.

### Multiple sequence alignment

MSA generated in ClustalW^51^ and represented in JalView^52^. The following TMEM263 sequences were used: *H. sapiens* isoform a (human): NP_001306590.1; *H. sapiens* iso b (human): NP_001306595.1; *P. macrocephalus* (sperm whale): XP_054941260.1; *D. rotundus* (common vampire bat): XP_045042869.1; *G. gallus* (chicken): NP_001006244.2; *L. disocolor* (Swift parrot): XP_065535720.1; *A. carolinensis* (green anole): XP_016849149.1; *C. mydas* (Green sea turtle): XP_037736208.1; *X. tropicalis* (tropical clawed frog): NP_989399.1; *M. unicolor* (Tiny Cayenne Caecilian): XP_030070213.1; *D. rerio* (zebrafish): NP_998306.1; *E. electricus* (electric eel): XP_026885649.1.

### AlphaFold2 modelling

Protein models were built using AlphaFold2 via ColabFold v1.5.2^53, 54^ using residues 33-116 of human TMEM263 taken from UniProt (Q8WUH6). To maximise the structural sampling, 8 independent seeds were run, to generate a total of 40 models. When ranked by pLDDT the top 5 models all had a similar double helix structure (Fig.3G) and an average RMSD to the top model of 0.69±0.33 Å. The AlphaFold model and all quality metrics are available to download on the Open Science Framework (https://osf.io/sfeur/).

**Figure 3.**
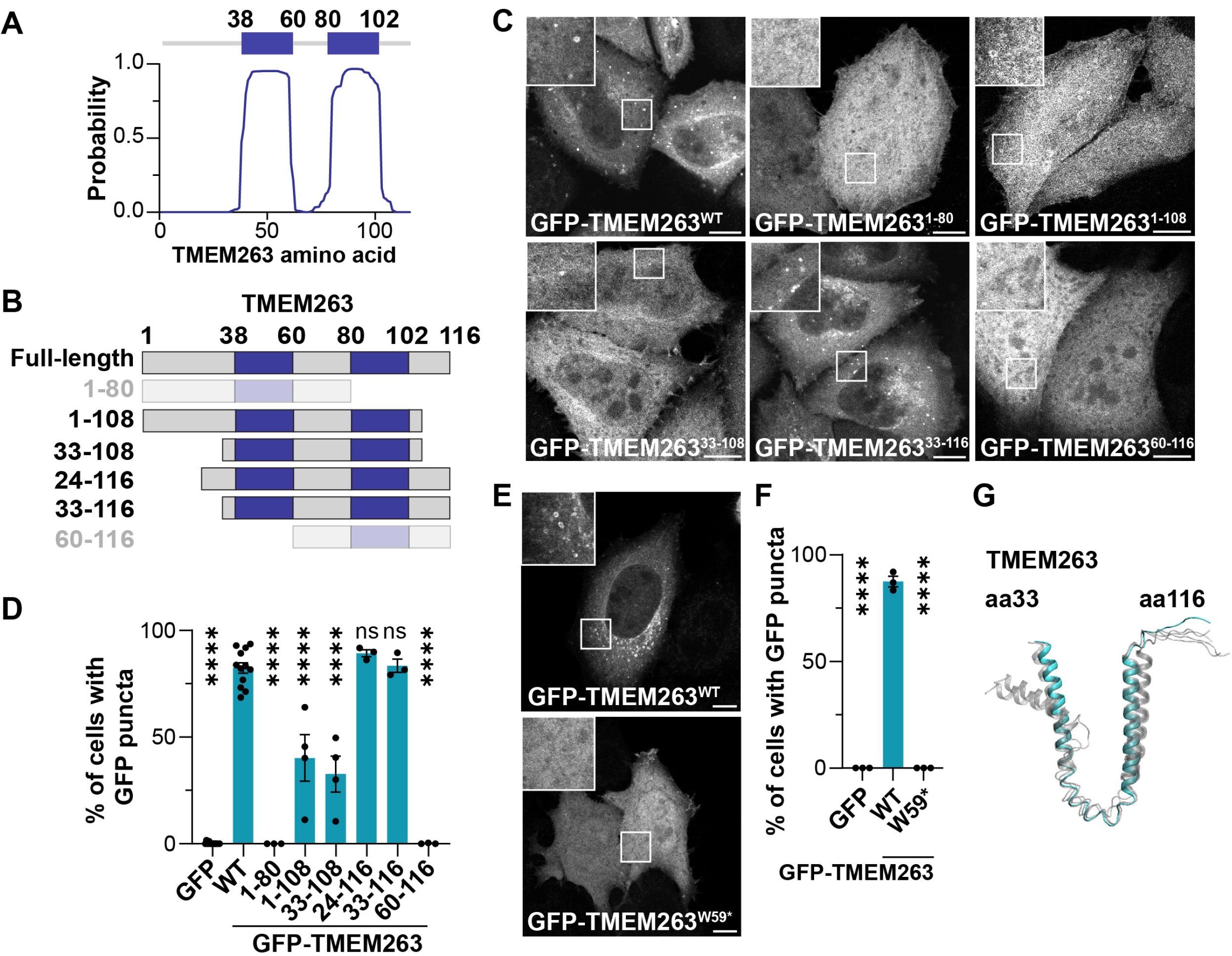
TMEM263 has features of a class I lipid droplet protein. **(A)** TMEM263 is an integral membrane protein. TMHMM predicts two transmembrane domains, amino acids 38-60 and 80-102 (labelled in dark blue) in *H. sapiens* TMEM263. **(B)** Diagram of TMEM263 protein truncations generated in this study. **(C,D)** TMEM263 transmembrane domains are required for its localisation. HeLa cells were transfected with GFP or GFP-TMEM263 truncations as indicated, were fixed and scored for the presence of GFP-positive puncta. **(E,F)** TMEM263 mutation associated with dwarfism impairs TMEM263 localisation. HeLa cells were transfected with GFP, GFP-TMEM263^WT^ or TMEM263^W59*^ as indicated, were fixed and scored for the presence of GFP-positive puncta. (G) TMEM263 folds into a hairpin structure. Overlay of the top 5 ranked of 40 AlphaFold2 models of TMEM263^33-116^. Rank 1 is coloured cyan and ranks 2-5 are coloured grey. Unless stated otherwise, n = 3. Scale bars are 10 µm. Bars extend to the SEM. For D and F, at least 600 cells per condition across n=3. Two-sided chi-squared tests were performed. ns p ≥ 0.05, **** p < 0.0001.

**Figure 4.**
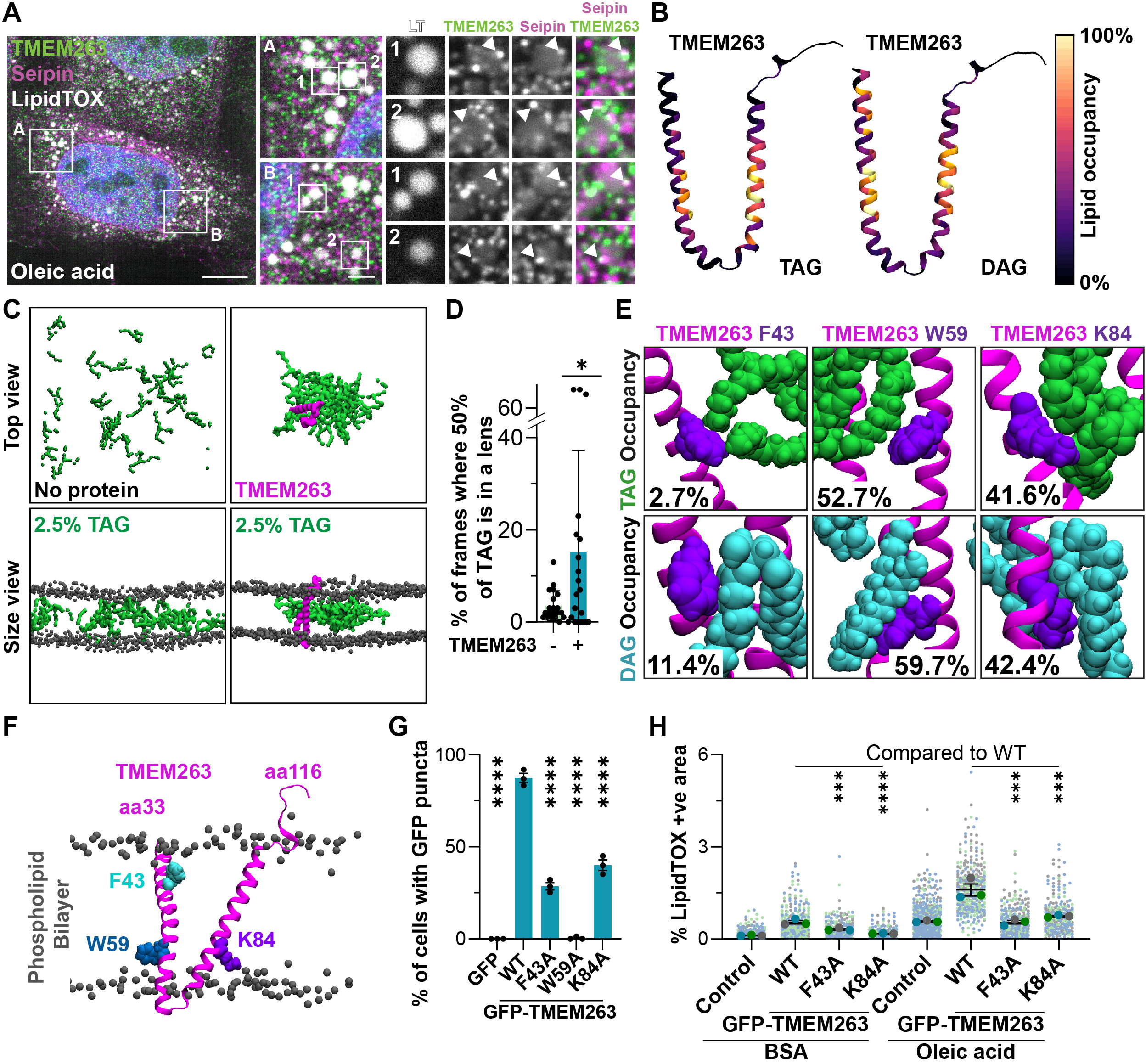
TMEM263 can bind and concentrate lipids in a phospholipid bilayer. **(A)** Endogenous TMEM263 co-occurs with lipid droplet biogenesis regulator, seipin. Representative images of HeLa cells treated with 30µM oleic acid for 16hours. Cells were fixed and immunostained with antibodies against endogenous TMEM263 and endogenous seipin, and stained with LipidTOX. White arrowheads highlights overlapping puncta. **(B)** PyLipID-derived TAG and DAG binding occupancies based on atomistic MD analysis. Occupancies were calculated for each residue as % of frames where the residue is within 0.35 nm of TAG/DAG. Data are projected as a coloured heatmap on the input AlphaFol2 model **C)** Representative frames of coarse-grained simulation of a bilayer containing 2.5% TAG with or without TMEM263^33-116^ present, viewed from the top and the side. **D)** Quantification of data from (C), measuring % of frames where at least 50% of TAG is present in the largest lens (cluster), with contacts within clusters defined as within 0.55nm. N=20 per condition. **E)** Representative images of simulations in (B) with F43, W59, and K84 displayed as spheres, and examples of TAG and DAG contacts shown. Occupancy values from PylipID analysis in B for these residues is listed. **F)** TMEM263^33-116^ (magenta) modelled in a bilayer during atomistic simulation (B) with residues F43, W59, and K84 as spheres. **(G,H)** Lipid binding residues are functionally important for TMEM263. HeLa cells were transfected with GFP or GFP-TMEM263 mutants as indicated, were fixed and scored for the presence of GFP-positive puncta **(G)**. At least 600 cells per condition total across n=3. Two-sided chi-squared tests were performed. Cells as in G were incubated for 16 hours with BSA vehicle control or 30µM oleic acid, fixed and stained with the neutral lipid dye, LipidTOX. The % LipidTOX positive area per cell was quantified **(H)**. Control represents untransfected cells in the same field of view as GFP-TMEM263-expressing cell. At least 200 cells per condition total across n=3. Ordinary one-way ANOVA with Dunnett’s multiple comparisons tests were performed. Data from control and GFP-TMEM263^WT^-expressing cells are also included in Fig. 2E. Scale bars are 10 µm, bars extend to the mean, colour coded individual data points correspond to individual cells from different experiments, lines show mean values, and error bars represent SEM. ns p ≥ 0.05, * p < 0.05, ** p < 0.01, *** p < 0.001, **** p < 0.0001.

**Figure 5.**
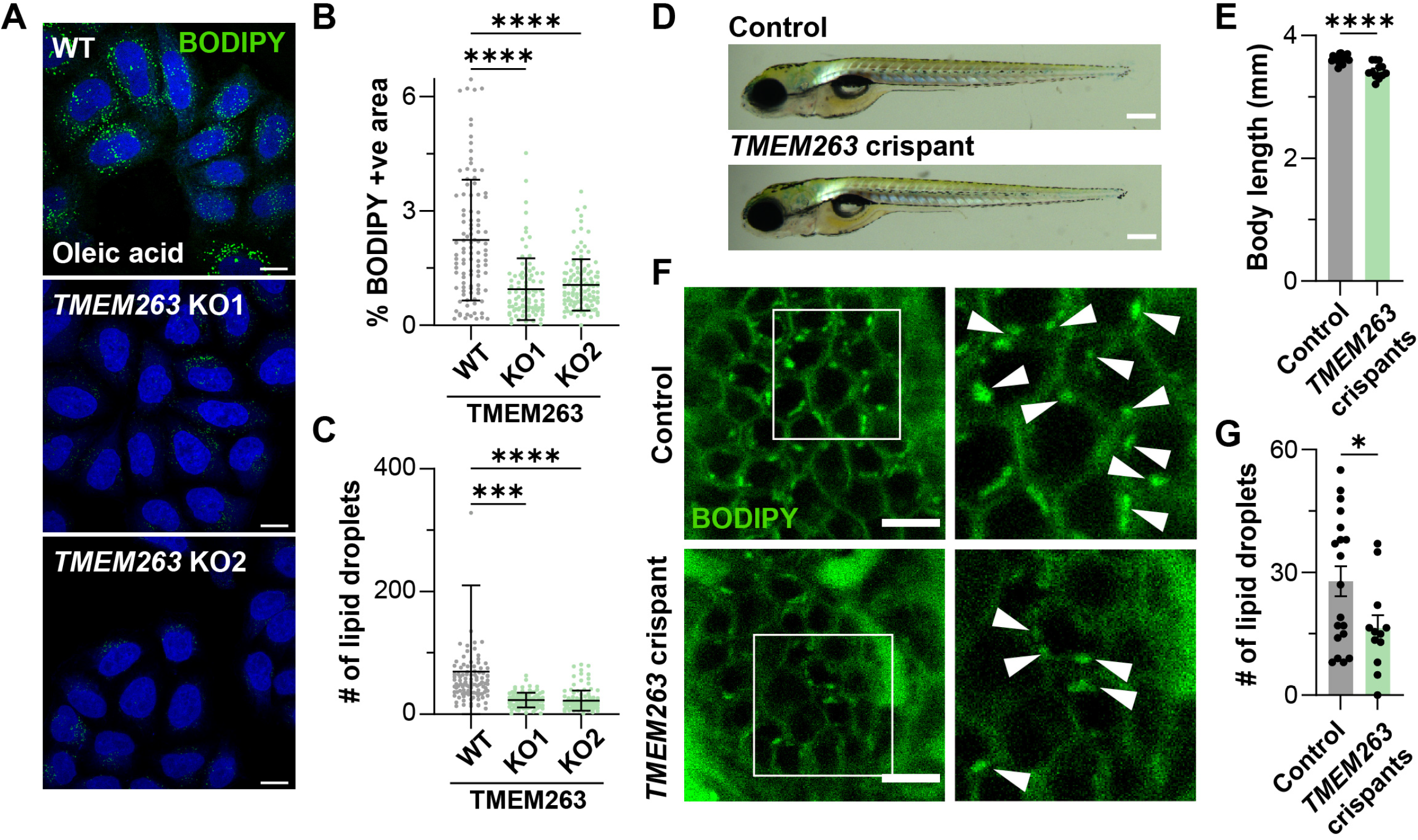
TMEM263 is required for efficient lipid droplet formation *in vivo*. **(A-C)** TMEM263 knock-out impairs lipid droplet formation. Parental, TMEM263 knock-out 1 (KO1) and TMEM263 KO2 HeLa cells were treated for 16 hours with 30µM oleic acid and stained with BODIPY **(A)**. The % BODIPY-positive area per cell **(B)** and number of lipid droplets per cell was quantified **(C)**. For B,C graphs show representative data from one repeat, n=3 with at least 90 cells per condition. Individual data points correspond to individual cells. Ordinary one-way ANOVA with Dunnett’s multiple comparisons test was performed. For C, 1 outlier is out of range of axis. **(D-G)** TMEM263 mutation in zebrafish impairs lipid droplet formation. **(D)** Representative images of wild-type control versus TMEM263 crispant zebrafish at 5 days post fertilisation (dpf). Scale bar = 250 μm. **(E)** Quantification of body length of control and TMEM263 crispant fish at 6dpf. *n* = 19 for *wt* and 12 for *tmem263;* Two-tailed unpaired student t-tests was performed. **(F)** Zebrafish (6dpf) were fed 300µM oleic acid for 24 hours, followed by incubation with BODIPY for 2 hours. Representative confocal microscope z-slices from intestine. Scale bars are 10 μm. White arrowheads in insets show lipid droplets. **(G)** Quantification of lipid droplet number per image from control and TMEM263 crispant fish. *n* = 19 for *wt* and 12 for *tmem263*. Student’s unpaired t-test was performed where * *p=*.0380. Lines show mean values, bars extend to the mean and error bars represent standard deviation. *** p < 0.001, **** p < 0.0001.

### Atomistic simulations

The AlphaFold2 TMEM263 model was used to seed molecular dynamic simulations. Models were built into simulation systems using CHARMM-GUI^55, 56^. Protein atoms were described with the CHARMM36m force field^57, 58^. Side chain pKas were assessed using propKa3.1^59^, and side chain side charge states were set to their default. The proteins were built into membranes comprising 4:4:2 DLPC, DMPC, and cholesterol. The shorter acyl tails of the DLPC/PE lipids were chosen to represent the thinner membrane of the ER. The membranes were solvated with TIP3P waters and neutralised with K^+^ and Cl^-^ to 150 mM. The simulation boxes were 8 × 8 × 8.5 nm and had ca. 55,000 atoms.

Each system was minimized and equilibrated according the standard CHARMM-GUI protocol, with a 50 ns final equilibration step. Production simulations were run in the NPT ensemble, with temperatures held at 303.5 K using a velocity-rescale thermostat and a coupling constant of 1 ps, and pressure maintained at 1 bar using a semi-isotropic Parrinello-Rahman pressure coupling with a coupling constant of 5 ps ^60, 61^. Short range van der Waals and electrostatics were cut-off at 1.2 nm. Simulations were run to 150 ns with 5 independent (reseeded) repeats. Lipid contacts were quantified using PyLipID^38^ implemented via LipIDens^62^.

### Coarse-grained simulations

Coarse-grained systems were built using the AlphaFold2 TMEM263 model. Systems were built using the Martini Maker package of CHARMM-GUI^63^. The proteins were described using Martini 2.2^64, 65^. Elastic networks of 500 kJ mol^-1^ nm^-2^ were applied between all protein backbone beads within 1 nm. All systems were solvated with Martini waters and Na^+^ and Cl^-^ ions to a neutral charge and 0.15 M. Minimization and equilibration followed the standard CHARMM-GUI Martini Maker protocol. Different systems were built, each described separately below.

The TAG blister systems were built using a monomer of TMEM263 in a membrane containing 2.5% TAG and 97.5% POPC. Boxes were set to 16 × 16 × 11.5 nm. Controls without protein were creating by deleting the protein beads from the system and equilibrating for 100 ns. Production simulations for each state were run for 2 µs with 20 repeats. TAG clusters were quantified using the GROMACS tool *gmx clustsize* with a cutoff of 0.55 nm for contacts. For each repeat, every frame was quantified for the total TAG that forms part of the main TAG cluster. Alternatively, a smaller system was built (12 × 12 × 12 nm) containing 59% POPC, 23% cholesterol, and 6% each of cholesterol ester (CE), DAG, and TAG. Post equilibration states were tiled into 9 repeats using the GROMACS tool gmx genconf and simulated for 7 µs to let TMEM263 interact with the CE/DAG/TAG blister. All simulations were run in GROMACS 2022.3^66^ [ref]. Data were analysed using Gromacs tools and VMD^67^.

### Other software utilised

TMHMM prediction was generated with TMHMM2.0^68^.

