## Supplementary figures and images for "The growth-promoting protein TMEM263 is an ER resident that controls early stages of lipid droplet biogenesis"

### Supplementary Figure S1-S5

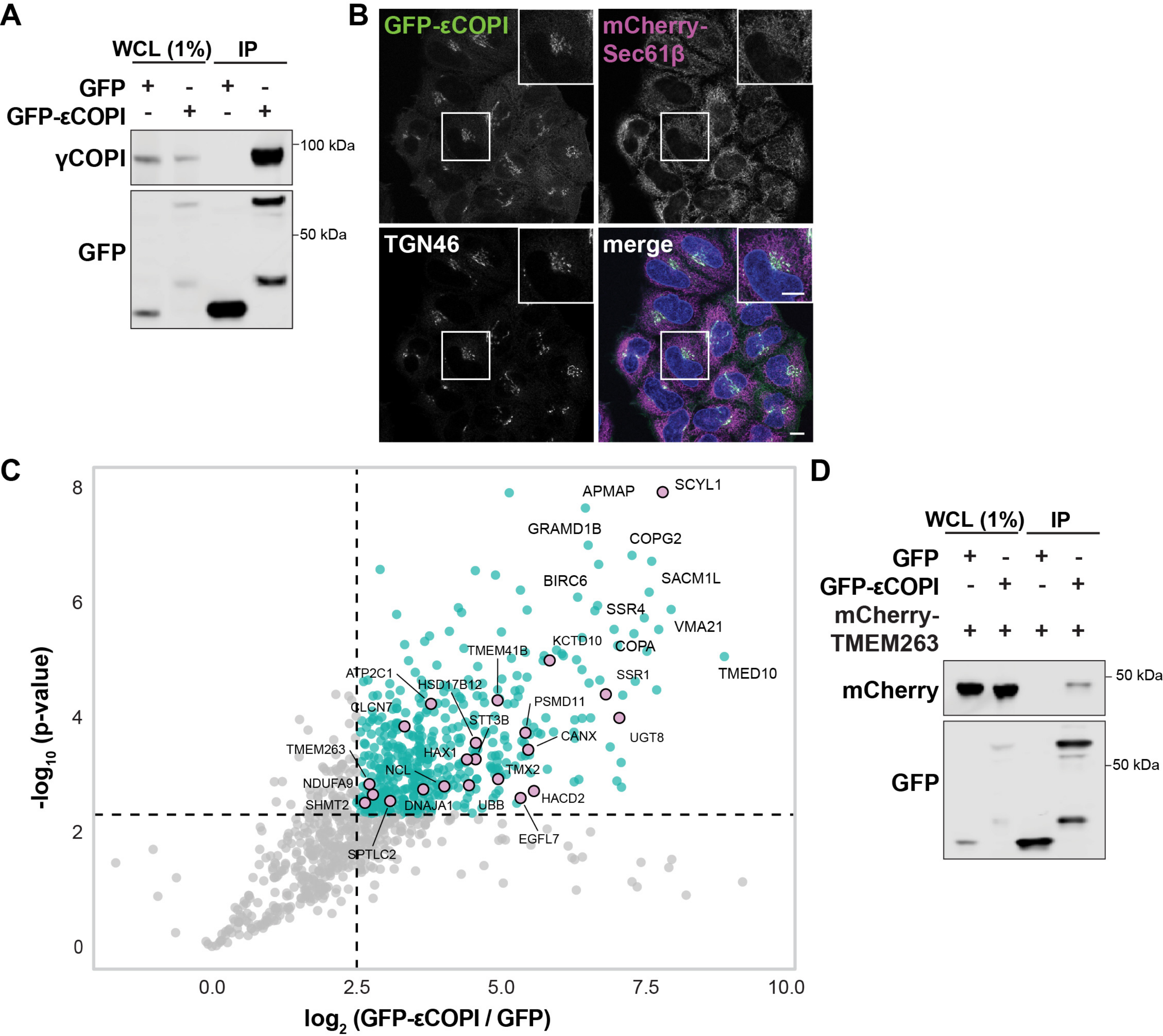

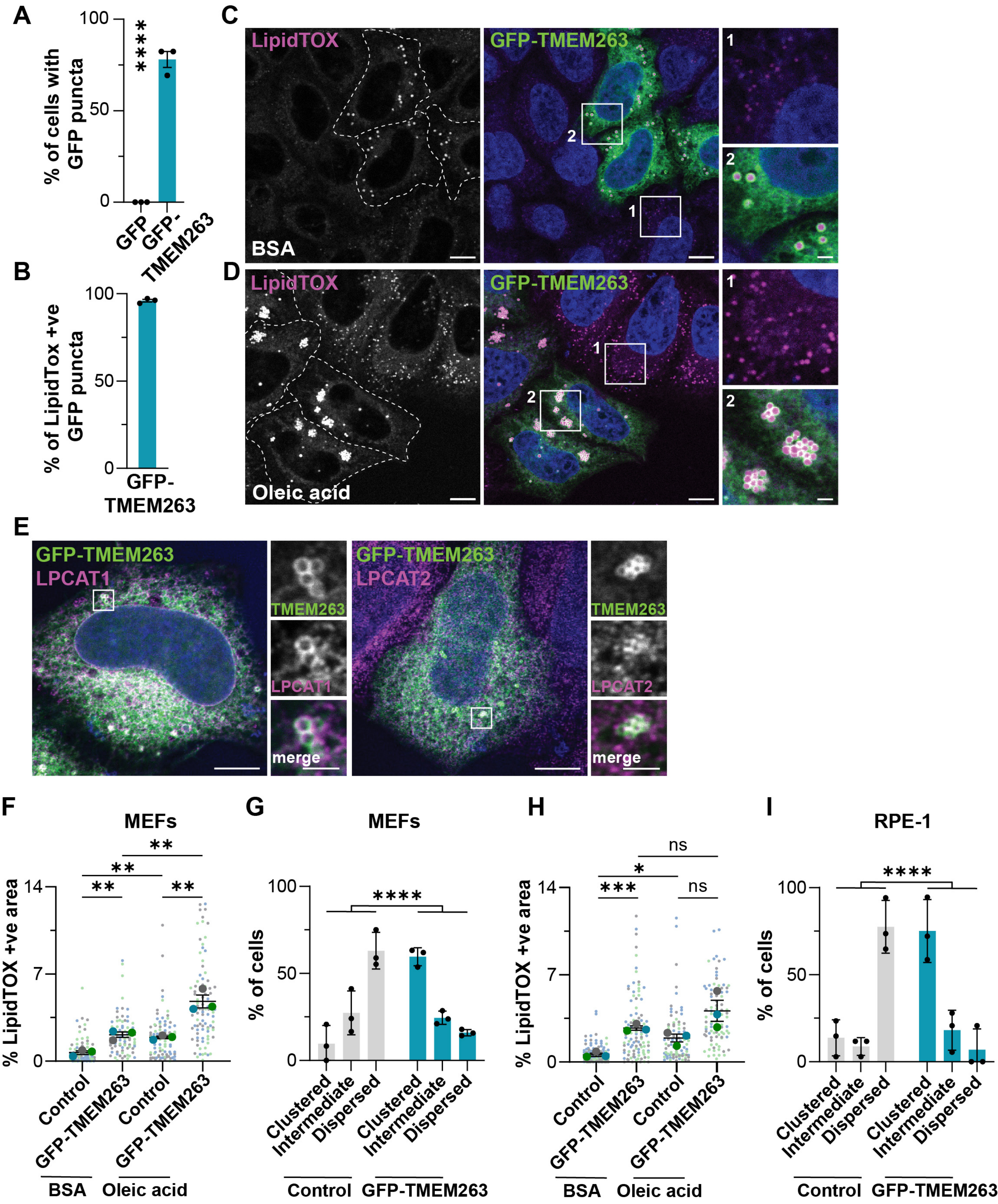

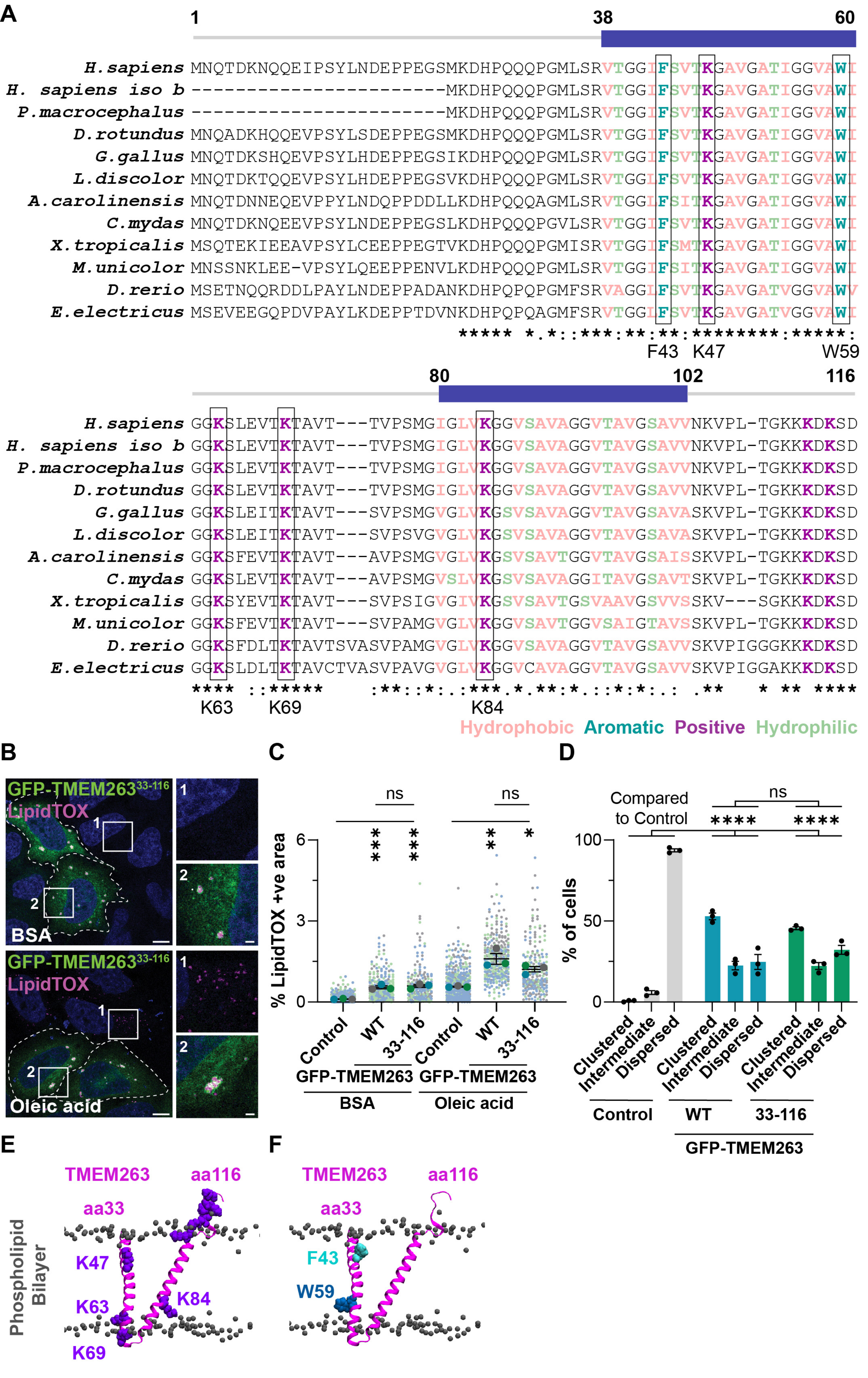

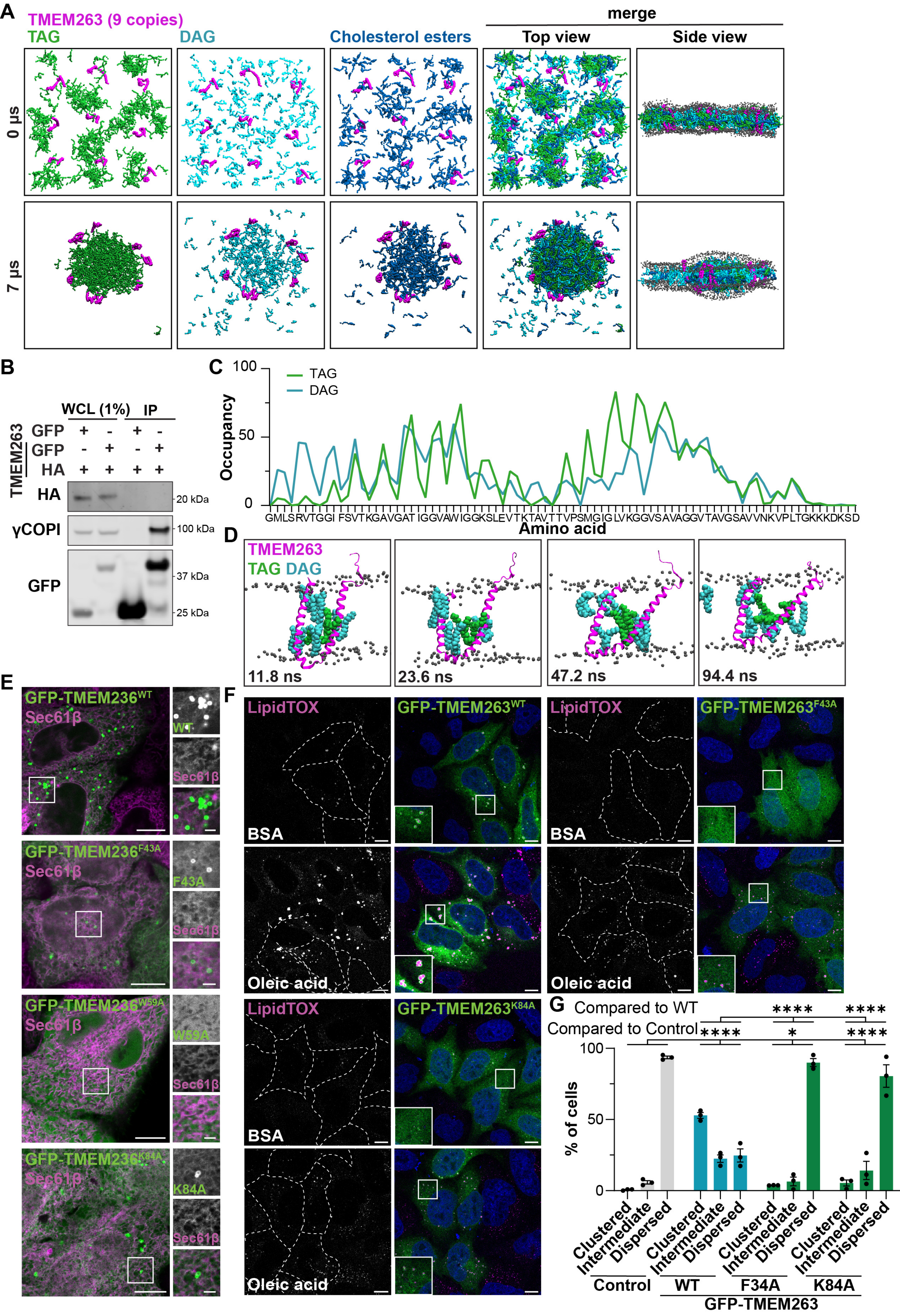

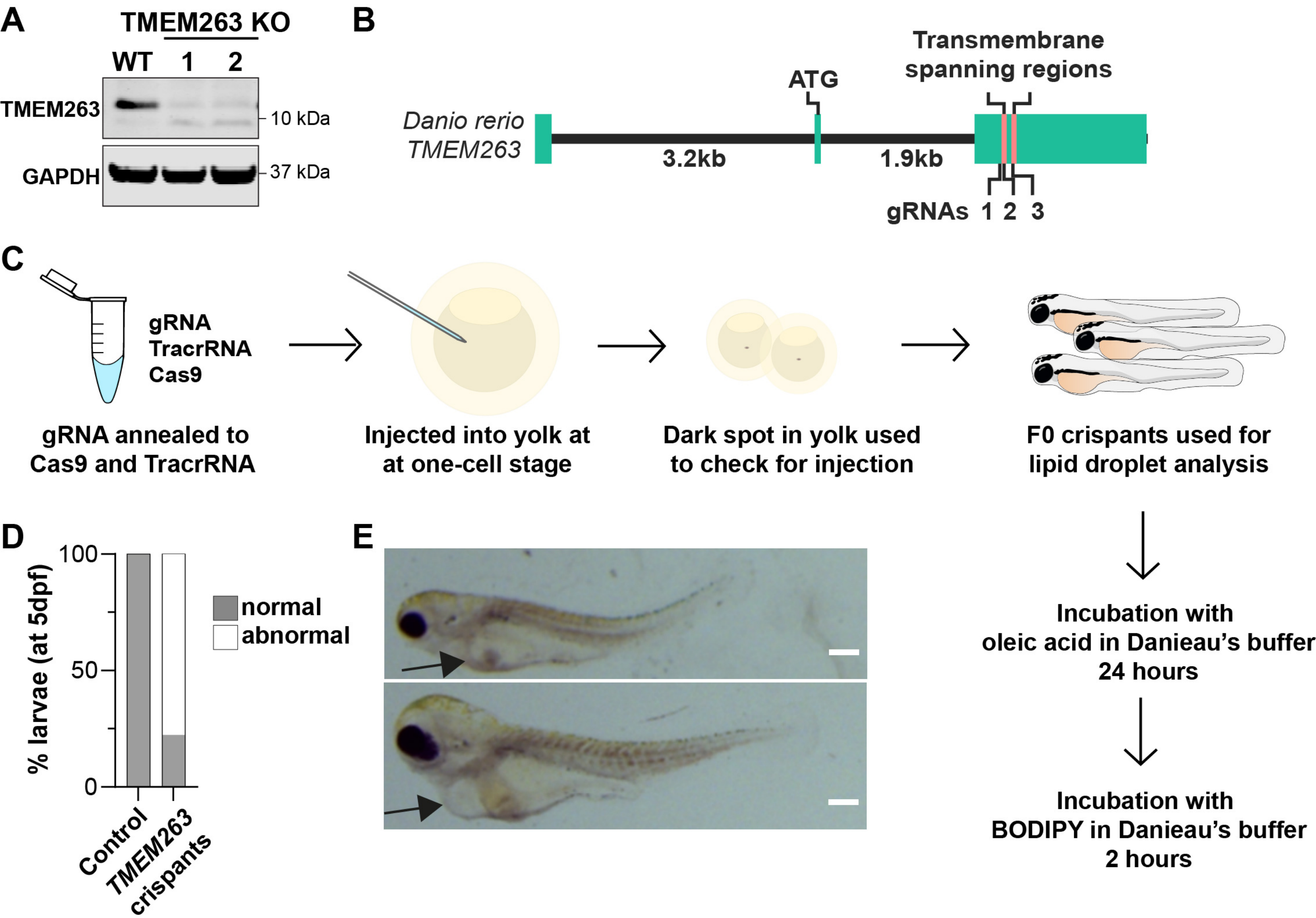
